# The Prokaryotic and Eukaryotic Microbiome of Pacific Oyster Spat is Shaped by Ocean Warming but not Acidification

**DOI:** 10.1101/2023.07.07.548145

**Authors:** Kevin Xu Zhong, Amy M. Chan, Brenna Collicutt, Maxim Daspe, Jan F. Finke, Megan Foss, Timothy J. Green, Christopher D.G. Harley, Amelia V. Hesketh, Kristina M. Miller, Sarah P. Otto, Kate Rolheiser, Rob Saunders, Ben J.G. Sutherland, Curtis A. Suttle

## Abstract

Pacific oysters (*Magallana gigas,* also known as *Crassostrea gigas*), the most widely farmed oysters, are under threat from climate change and emerging pathogens. In part, their resilience may be affected by their microbiome, which, in turn, may be influenced by ocean warming and acidification. Consequently, for three weeks, we exposed early-development Pacific oyster spat to different temperatures (18 and 24 °C) and *p*CO_2_ levels (800, 1600 and 2800 *µ*atm) in a fully crossed design. Under all conditions, the microbiome developed over time, with potentially pathogenic ciliates (*Uronema marinum*) greatly reduced in all treatments, suggesting that the spat’s microbiome undergoes adaptive shifts as the oysters age. The microbiome composition also differed significantly with temperature, but not acidification, indicating that *M. gigas* spat microbiomes can be altered by ocean warming but resilient to ocean acidification in our experiments. These findings highlight the spat microbiome’s flexibility to environmental changes as well as its “protective” capability against potentially pathogenic microbes.

## Introduction

The Pacific oyster (*Magallana gigas,* formerly known as *Crassostrea gigas*) is the most economically important and widely farmed species of oyster (Botta et al. 2020), and as such, serves as a model organism for examining the influence of environmental change on bivalves (Li et al. 2021; Li et al. 2018; Zhang et al. 2012). These oysters, like many other bivalves, are under threat from increasing ocean acidification and warming from climate change (Hoegh-Guldberg and Bruno 2010; Lokmer and Mathias Wegner 2015; Soon and Zheng 2020), as well as the emergence of pathogens (Altizer et al. 2013; Lokmer and Mathias Wegner 2015; Rowley et al. 2014; Zgouridou et al. 2022). One life stage that is particularly susceptible to die-offs is oysters that have recently undergone metamorphosis from pelagic larvae to benthic juveniles (spat) (Foulon et al. 2018). Such metamorphosis is irreversible and essential for survival, but is also a stage in which mortality and morbidity are often reported (King et al. 2019b).

The microbiome comprises bacteria, archaea, microeukaryotes (protists and fungi) and viruses that associate with a multicellular host, and can influence host health, biology, and performance across a variety of organisms (Apprill 2017; Knight et al. 2017; McFall-Ngai et al. 2013), including oysters (King et al. 2019b). In oysters, the microbiome includes the planktonic microbes on which it feeds, such as phytoplankton (Dupuy et al. 1999; Kamiyama 2011; Weissberger and Glibert 2021), as well as all the microbes associated with the oyster. This microbial consortium can have positive and negative effects on oyster growth and health. For example, the microbiota within the oyster digestive system can contribute to digestion (King et al. 2012) and nutrient uptake (Vogeler et al. 2022). In addition, microbes can promote oyster settlement and metamorphosis (Dobretsov and Rittschof 2020; Yu et al. 2010; Zhao et al. 2003) and assist in defense against infection by outcompeting pathogenic bacteria, producing antimicrobial compounds, and stimulating the host’s immune system (Desriac et al. 2014; King et al. 2019b; Lokmer and Mathias Wegner 2015). Yet, microbes can also cause disease in larval, juvenile, and adult oysters, leading to devastating mortality in oyster hatcheries and farms (King et al. 2019b). Putative microbial pathogens include bacteria (e.g., *Vibrio* spp., *Nocardia crassostreae*, *Roseovarius crassostreae*; Boardman et al. 2008; Friedman et al. 1991; Jeffries 1982), protists (e.g., *Mikrocytos mackini*, *Uronema marinum*, *Perkinsus marinus*; Andrews 1988; Bower 2010; Elston et al. 2015; King et al. 2019b; Plunket and Hidu 1978), fungi (e.g., *Ostracoblable implexa*; Bower 2010), viruses (e.g., OsHV-1; Renault et al. 2014), and even algae (Landsberg 2002). Microbe-mediated oyster mortality seems to be of polymicrobial origin (Dupont et al. 2020) and can be affected by several abiotic factors, such as temperature, pH, salinity, and nutrients (King et al. 2019b). For example, in oysters, elevated temperature can trigger disease outbreaks associated with *Vibrio* spp. (Green et al. 2019) and *Perkinus marinus* (Cook et al. 1998).

Oyster disease and mortality are strongly associated with higher temperatures and ocean acidification, both of which are exacerbated by climate change. Indeed, in some areas, anthropogenic greenhouse gas emissions (e.g., CO_2_) are predicted to lead to global surface temperature increases of up to 5.4°C (IPCC 2014; Tollefson 2020) and warming of ocean surface waters by 1.5°C (Bindoff et al. 2019) by the year 2100. As well, the partial pressure of carbon dioxide (*p*CO_2_) in seawater is expected to rise from about 400 to 1000 ppm, amounting to a 0.29 decrease in pH (Bindoff et al. 2019). Superimposed on these overall trends is local variability, which can lead to more rapid short-term changes. For example, this study was based in the Strait of Georgia, an area with large-scale production of Pacific oysters. Here, changes in atmospheric forcing and ocean circulation were associated with a warming trend of 0.024°C y^-1^ throughout the water column; overlaid on this was interannual variability (Masson and Cummins 2007). While variability exists in the carbonate chemistry of the Salish Sea, several studies have reported instances where the pH of the waters can reach relatively low levels for seawater (∼7.8), and the water column is undersaturated with respect to aragonite (Ianson et al., 2016; Evans et al. 2019; Hare et al. 2020), conditions that can be detrimental to shellfish survival (Haigh et al. 2015). Moreover, strong northwesterly winds can cause short-lived events of elevated *p*CO_2_ and low-pH, combined with corrosive aragonite conditions in the northern Strait of Georgia (Evans et al. 2019). Consequently, short-term excursions of low pH waters are becoming increasingly common under a scenario of climate change. In turn, increasing acidity and temperature can profoundly affect marine ecosystems (Hoegh-Guldberg and Bruno 2010), including influencing the microbiome of oysters (Fernandez-Piquer et al. 2012; Lim et al. 2021a; Lokmer and Mathias Wegner 2015; Scanes et al. 2021a; Scanes et al. 2021b; Scanes et al. 2021c; Unzueta-Martínez et al. 2021) and the interaction of the oyster immune system with the microbiome (Dang et al. 2023). Despite the potential influence of the microbiome on oyster health, and the frailty of spat to environmental change, the impact of temperature and *p*CO_2_ on oyster spat remains unexplored, as does effects on the eukaryotic microbiome of oysters, in general. Understanding the response of spat and their microbiome to these stressors can inform hatchery practices, enhance the resilience of the spat prior to their deployment, and improve our knowledge of their survival and growth performance in the changing natural environment.

Given the critical role that bivalves play in ecosystem function, the importance of *M. gigas* as a model organism and aquaculture species, and the precarious position of calcifying organisms in the face of rising temperatures and ocean acidification, we conducted a fully crossed experiment to examine the impact of different temperatures and *p*CO_2_ levels on the microbiome of oyster spat. For three weeks, we followed changes in the prokaryotic and eukaryotic microbiome of oyster spat grown at 18°C and 24°C, and *p*CO_2_ levels of 800, 1600 and 2800 *µ*atm. Temperature and maturation, but not acidity, were associated with significant changes in the microbiome. As well, a small number of taxa comprised a core microbiome. These data provide insight into how the microbiome of oyster spat responds to episodic changes in the temperature and acidity of seawater, which are expected to increase in frequency as the result of climate change.

## Materials and Methods

### Experimental materials and procedures

During the summer of 2019 (July 24 to August 16) at the Hakai Institute located on Quadra Island (BC, Canada), we conducted a challenge study that incubated oyster spat in 340-L mesocosm tanks under six different computer-controlled temperature and *p*CO_2_ treatments (18°C+800*µ*atm, 18°C+1600*µ*atm, 18°C+2800*µ*atm, 24°C+800*µ*atm, 24°C+1600*µ*atm, 24°C+2800*µ*atm) with 5 replicate tanks for each treatment. The 18°C+800*µ*atm treatment served as a “control” treatment, which is similar to the local environmental conditions experienced by Pacific oyster spat (Ianson et al., 2016; Evans et al. 2019; Hare et al. 2020; Haigh et al. 2015; Evans et al. 2022). In each tank, water temperatures were maintained by circulating hot or cold glycol through a titanium heat exchange coil, controlled by a Dwyer TSW-250 Dual Stage Temperature Switch (Dwyer Instruments Inc). The *p*CO_2_ conditions were achieved by mixing 99% carbon dioxide gas and ambient air through mass flow controllers (Model MC-500SCCM-D-DB9S and MCR-1000SLPM-D-DB9S; Alicat Scientific). Treatment gas mixtures were bubbled into mesocosms with an airstone. The monitoring of temperature and pH of water in the tank was performed using Walchem 600 Series Model WCT600HSNNE-NN with Omega HSRTD-3-100-A-120-E-ROHs temperature probes and Honeywell Durafet III pH electrodes. At every sampling time point, discrete water samples were collected in 350 mL amber glass bottles to constrain the full carbonate chemistry and to measure and validate the *p*CO_2_ values using a Burke-o-Lator containing a LI-COR non-dispersive infrared absorbance gas analyzer (L1840A; LI-COR Biosciences), following the methods described in Evans et al. (2019) and (2022). Total carbon dioxide (TCO_2_) concentrations were measured and validated using certified reference materials (CRM) provided by A. Dickson (Scripps Institute of Oceanography), while *p*CO_2_ measurements were validated with certified gas standards. Both measurements had analytical uncertainty values of equal or less than 1% (Evans et al. 2019; Evans et al. 2022). These measurements were also used to calculate the average pH achieved for each treatment over the course of the challenge experiment. Data was processed using an Excel version of CO2SYS version 2.1 (Pierrot et al. 2011) (carbonic acid dissociation constants form (Lueker et al. 2000), bisulphate dissociation constant form (Dickson et al. 1990), and the boron/chlorinity ratio form (Uppstrom 1974)).

For the challenge study, a cohort of oyster spat (∼500 µm in shell height), were produced in-house at the Deep Bay Marine Field Station of Vancouver Island University from a select group of adult Pacific oysters known for their robustness and high survival rates. The spat were produced from a cross between four Chilean females with six USA males. To obtain a diverse array of microbes, each mesocosm was filled with ∼340 L of untreated seawater (salinity = ∼29 PSU) comprised of 15% deeply-mixed water from Seymour Narrows (50.1372° N, 125.3537° W) and 85% near-surface water from stratified Hyacinthe Bay (50.1220° N, 125.2261° W). For each of the 30 mesocosms (n = 5 tanks per treatment), ∼33,000 individual ∼0.5-mm oyster spat were added to a “condo” (21.5-cm in diameter and 15-cm in height) made of PVC pipe and with 250-µm mesh Nitex screening glued to each end to allow water and food to pass. Tank water was circulated upwards through the pipe to create a gentle upwelling effect. Oysters were acclimated in tanks for 24 h at 18°C and 800 *µ*atm *p*CO_2_ (T0). After acclimation, the temperature and *p*CO_2_ levels in the tanks were gradually adjusted over the course of 48 h, so that half of the tanks were at 24°C while half remained at 18°C, and five tanks at each temperature had *p*CO_2_ levels of 800 *µ*atm, 1600 *µ*atm, and 2800 *µ*atm. The average pH achieved over the course of the experiment for each treatment (calculated pH mean ± SD, n = 30) was 7.79 ± 0.09, 7.50 ± 0.04, 7.24 ± 0.04, 7.76 ± 0.08, 7.48 ± 0.05 and 7.28 ± 0.04 for 18°C+800µatm, 18°C+1600µatm, 18°C+2800µatm, 24°C+800µatm, 24°C+1600µatm and 24°C+2800µatm treatments, respectively (Table S1). Once temperature and *p*CO_2_ conditions were stabilized after 48 h, the experiment began (T1). Additionally, ∼20% of mesocosm water was replaced each day by a continuous flow of 50 mL min^-1^ of 1-µm filtered and UV-treated seawater from Hyacinthe Bay. Oysters were statically fed each day for two hours at a concentration of 50,000 cells mL^-1^ consisting of 50% *Tisochrysis lutea* paste (Instant Algae ISO 1800) and 50% live cultured *Chaetoceros muelleri*.

Every three to four days, for three weeks, duplicate samples of ∼50 to 100 spat (for microbiome analysis) were collected from each tank, prior to feeding, resulting in six sampling time points (T1 to T6); as well, samples were collected at T-1 and T0. During sampling, the oyster “condo” (a structure used to house the oysters during the experiment) was removed from the tank and placed in a clean plastic bin. Oyster samples were collected using a sterile spatula, placed in cryovials, immediately frozen using liquid nitrogen, and stored at −80 °C until processed for microbiome analysis. Notably, for each treatment, we utilized samples from three out of the five replicate tanks (replicate: A, B, and C) for the purpose of microbiome analysis in this study.

### Genomic DNA extraction

For each sample, ∼50 individuals were homogenized in a 1.5-mL microcentrifuge tube using a sterile Tissue Grinder (Fisher Scientific). Genomic DNA was then isolated using ZymoBIOMICS DNA Miniprep Kits (Zymo Research) following the manufacturer’s instructions. DNA was stored at −80°C prior to further analysis.

### Deep sequencing of 16S and 18S rRNA genes

Deep sequencing of 16S and 18S rRNA gene amplicons was employed to profile the prokaryotic and eukaryotic microbiota, respectively, associated with the oysters.

The preparation of 16S amplicon sequencing libraries was adapted from the online Illumina protocol (Amplicon et al. 2013) using 515F-Nxt and 806R-Nxt primers (Apprill et al. 2015) that targeted a ∼292-bp 16S rRNA gene fragment in the V4 region (Table S2). See the Supplementary Methods for details.

To prepare the 18S amplicon sequencing library for microeukaryotes, we employed the CRISPR-Cas Selective Amplicon Sequencing (CCSAS), a method developed by Zhong et al. (2021). Briefly, following amplification with the universal primers TAReuk454FWD1-Nxt and TAReukREV3-Nxt (Table S2) (Stoeck et al. 2010; Zhong et al. 2021), CRISPR-Cas combined with the oyster-specific gRNA m258 (the 20-nt gRNA-target-site oligo sequence: 5’-AGTAAACGCTTCGGACCCCG-3’) was used to cut the oyster 18S rRNA gene amplicons, followed by DNA size-selection using SPRIselect beads (Beckman Coulter) to obtain the intact amplicons to produce a host-18S reduced amplicon sequencing library.

The 16S and 18S amplicon sequencing libraries were quantified using a Qubit® dsDNA HS Assay Kit (Invitrogen). The fragment size was then determined using the Agilent High Sensitivity DNA Kit on an Agilent 2100 Bioanalyzer System. Equimolar amounts of these barcoded amplicon-sequencing libraries were pooled and sent to the University of British Columbia’s BRC-Seq Next-Gen Sequencing Core for sequencing using the MiSeq platform with 300-bp pair-end chemistry (Illumina).

### Amplicon sequence analysis

Amplicon sequences for both 16S and 18S rRNA genes were processed and analyzed using QIIME pipeline version 2 (qiime2.2019.04) (Caporaso et al. 2010). Briefly, adapters and low-quality reads were trimmed with Trimmomatic v.0.36 (Bolger et al. 2014) using the following settings (EADING:3 TRAILING:3 SLIDINGWINDOW:4:15 MINLEN:36). Paired-end reads were then merged using PEAR (Zhang et al. 2014) with the default settings. These assembled sequences were then loaded into QIIME v.2 (Caporaso et al. 2010), and DADA2 (Callahan et al. 2016) was used to filter noisy and chimeric sequences, allowing us to obtain amplicon sequence variant (ASV) features.

Taxonomic position was assigned for these ASV features using a pre-built Naïve Bayes classifier that was trained on the SILVA v.132 16S or 18S SSU database (Quast et al. 2013) at 99% nucleotide sequence similarity for 16S and 18S rRNA genes, respectively. ASV features for 16S rRNA genes were removed from downstream analysis for those with taxonomy-assignment confidence less than 80%, if unclassified at the kingdom level, or belonging to eukaryotes, mitochondria, and chloroplasts. Features for 18S rRNA genes were removed for those with taxonomy-assignment confidence less than 90%, if unclassified at the kingdom level, or belonging to bacteria, archaea, mitochondria, and chloroplasts.

For both 16S and 18S rRNA genes, to minimize stochastic effects resulting from rare ASVs, we set two reads as the cutoff value for the minimum number of reads for analysis: the number of reads ≥2 was considered and kept; otherwise, the value was set as zero. To produce representative ribosomal gene communities per library, the ASV feature table was normalized to the lowest number of total reads per library by rarefication using Phyloseq v.1.26.1 (McMurdie and Holmes 2013) in R v.4.0.3 (R Core Team 2013). The downstream analysis was conducted using R packages Phyloseq v.1.26.1 (McMurdie and Holmes 2013), Microbiome v.1.13.12 (Lahti and Shetty 2018), and Vegan v.2.5 (Oksanen 2013), with figures generated using ggplot2-v3.3.3 (Wickham 2016).

### Statistical analysis

Indices of alpha-diversity (e.g., the observed ASV richness, Pielou’s evenness index (*J’)*, and Shannon’s diversity index (*H’*)) and beta-diversity (e.g., Pearson correlation coefficient, Weighted unifrac distance matric) were calculated using Phyloseq v.1.26.1 (McMurdie and Holmes 2013), microeco v.0.11.0 (Liu et al. 2021), Vegan v.2.5 (Oksanen 2013), and QIIME v.2 (Caporaso et al. 2010). The influence of temperature, *p*CO_2_ and time on alpha-diversity, either individually or in combination, was examined using ANOVA (with temperature, *p*CO_2_ and time included in the case of three-way ANOVA). Post-hoc analysis was then conducted using Tukey’s Honest Significant Difference (HSD) test, facilitated by the R stats package.

To compare prokaryotic and microeukaryotic community structures among samples, principal coordinate analyses (PCoA) were performed based on Bray-Curtis dissimilarities in the ordination of the weighted UniFrac metrics (Lozupone et al. 2011), which consider both the presence/absence and the relative abundance of ASVs. The dissimilarity of the prokaryotic and microeukaryotic community composition as a function of temperature, *p*CO_2_, and time, either individually or in combination, was examined using PERMANOVA (Anderson 2014) with the adonis function and Bray-Curtis method in Vegan v.2.5 (Oksanen 2013). The pairwise post-hoc test was performed using pairwise-Adonis v.0.4 in R (Martinez Arbizu 2017). Pearson correlation coefficients of prokaryotic and microeukaryotic community structure across samples were visualized using corrplot v.0.90 (Wei et al. 2017).

Differences in the core microbiome of oyster spat among treatments and time points were examined using R package Microbiome v.1.13.12 (Lahti and Shetty 2018). As followed in previous studies (Miller et al. 2020; Unzueta-Martínez et al. 2022) and based on our sample size and sequencing depth (Neu et al. 2021), ASVs with a frequency of occurrence >50% across all examined samples (e.g., by treatments or time-points) and a relative abundance >1% were considered as core constituents of the microbiome of oysters.

Linear discriminant analysis Effect Size (LEfSe) (Segata et al. 2012) was conducted to identify prokaryotic and microeukaryotic taxa that were differentially abundant among treatments and time points and were calculated using default settings (*alpha* value cutoff for the factorial Kruskal-Wallis test among classes is 0.05; threshold on the logarithmic LDA score for discriminative features is 2.0) of the Galaxy modules (Afgan et al. 2018) provided by the Huttenhower lab.

To investigate the relationships between microbial community structure and environmental variables (temperature, *p*CO_2_, and pH), a Canonical Correspondence Analysis (CCA) was performed using Vegan v.2.5 (Oksanen 2013). Variables were tested for significance using a forward selection procedure and an unrestricted Monte Carlo permutation test (999 permutations).

We also computed the pairwise Euclidean distance between the samples for temperature, *p*CO_2_, pH, time (days since the experiment started), and the prokaryotic and microeukaryotic alpha-diversity indices using Pearson correlation analysis using Vegan v.2.5 (Oksanen 2013). Then, the partial Mantel test was performed to assess the pairwise correlation between the prokaryotic and microeukaryotic community composition and the measured variables, respectively.

## Screening for potential pathogen species

To identify potential pathogen species associated with Pacific oyster spat, we screened both the prokaryotic and microeukaryotic ASVs against Bower’s list of pathogen species in shellfish (Bower 2010) using the ‘grepl’ function in R. ASVs were selected if their taxonomy assignment at the species level matched the name of a recognized pathogen.

## Results

### Variability of prokaryotic communities in oyster spat across time and treatments

Deep amplicon-sequencing, which targets a 292-bp fragment of the 16S rRNA gene, was used to profile the oyster-associated prokaryotic community. A total of 14,231 prokaryotic ASVs from 16 bacterial and three archaeal phyla were discovered in oyster samples across treatments and time points (Fig. 1a; Fig. S1). Overall, members of the bacterial phyla Bacteroidetes, Proteobacteria, and GN02 (a.k.a. Gracilibacteria) dominated the oyster spat microbiome at the beginning of the experiment (T-1) and following 24-hours of acclimation (T0). However, a decrease of bacteria in GN02, accompanied by enrichment of those in the Chloroflexi, Planctomycetes, Actinobacteria, TM6, and Verrucomicrobia occurred across temperatures, *p*CO_2_ treatments and time.

**Fig. 1.**
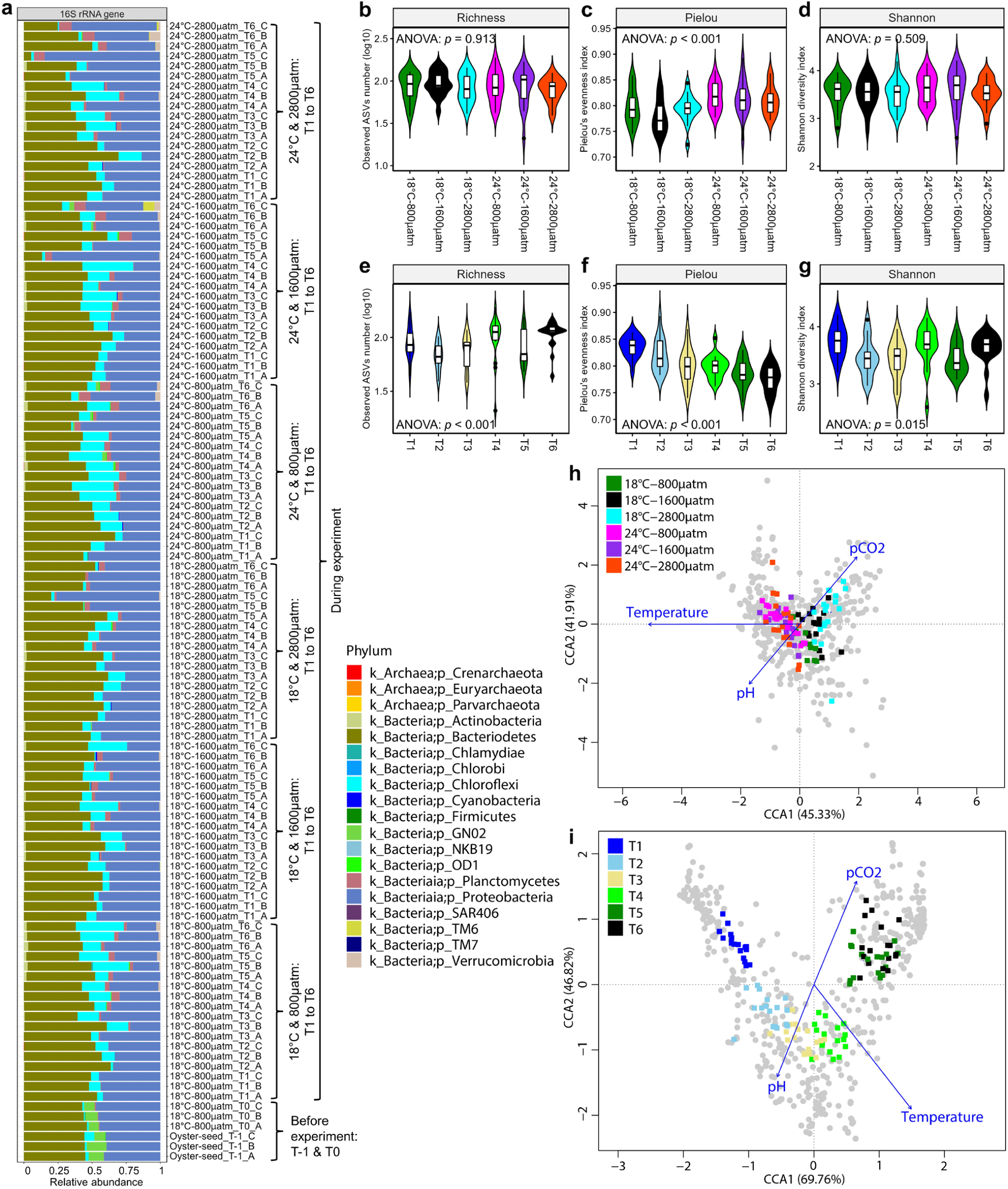
Variability of prokaryotic community associated with Pacific oyster spat during the challenge experiment. **a** Relative abundance of prokaryotic 16S rRNA gene sequences in oysters spat samples subjected to different temperature (18℃, 24℃) and *p*CO_2_ (800, 1600, and 2800 µatm) conditions over the course of three weeks (T1 to T6). Additionally, oyster spat samples before the experiment (T-1: original oyster spat; T0: oyster spat sampled after 24-hour acclimation in tank-water before the experiment began) were also examined. **b-g** Alpha diversity (e.g., richness, Pielou’s *J*, and Shannon’s *H*) of prokaryotic 16S rRNA genes across time and treatments. The violin plot is a smoothed density plot of the data, with the violin’s width representing the frequency of observations in that range of values. The line in the middle of the violin plot represents the data’s median, and the square indicates the data’s mean. **h & i** Canonical Correspondence Analysis (CCA) with measured abiotic variables within mesocosms (temperature, *p*CO_2_, and pH) as constrained variables to explain community changes of prokaryotic 16S rRNA gene sequences associated with oysters between treatments (**h**) and time points (**i**). Grey points represent different prokaryotic ASVs, and the square shapes indicate the oyster samples. Panel **b-d** shares the same color code as panel **h**, while panel **e-g** shares the same color code as panel **i**.

We measured alpha-diversity across time points and treatments and observed that richness ranged between 21 and 189, averaging 91 ± 33 (mean ± SD, n = 108; Fig. 1b,e), Pielou’s evenness index (*J’*) varied between 0.72 and 0.89, with an average of 0.80 ± 0.03 (mean ± SD, n = 108; Fig. 1c,f), and the Shannon diversity index (*H’*) ranged between 2.59 and 4.21, with an average of 3.56 ± 0.33 (mean ± SD, n = 108; Fig. 1d,g). Significant differences in richness (ANOVA: *p* < 0.001), Pielou’s *J* (ANOVA: *p* < 0.001), and Shannon’s *H* (ANOVA: *p* = 0.015) occurred among six-time points (Fig. 1e-g). However, neither temperature nor *p*CO_2_ significantly affected richness or Shannon’s *H* (Fig. 1b,d; Fig. S2). Elevated temperatures resulted in significantly higher community evenness under low and intermediate, but not high, *p*CO_2_ conditions (ANOVA: *p* < 0.001; Fig. 1c).

The prokaryotic community structure varied with time and treatment (Fig. S1; Fig. S3-4; Fig. 1). Non-significant difference in microbiome composition was detected among the six combinations of temperature and *p*CO_2_ (PERMANOVA: r^2^ = 0.054, *p* = 0.066), or among the three *p*CO_2_ levels (PERMANOVA: r^2^ = 0.019, *p* = 0.386). However, the variation was significant between temperatures (PERMANOVA: r^2^ = 0.022, *p* = 0.002) and among the six-time points (T1 to T6; PERMANOVA: r^2^ = 0.199, *p* = 0.001). These results are supported by PCoA and CCA, which showed that prokaryotic communities at similar temperatures (18℃, 24℃) and time points (T1 to T6) roughly clustered together (Fig. 1h-i; Fig. S4).

### Variability of microeukaryotic community in oyster spat across time and treatments

The taxonomic distribution of 18S rRNA genes of microeukaryotes associated with oysters was obtained using amplicon deep sequencing. A total of 36,499 microeukaryotic ASVs from 19 fungal and 180 protistan genera were detected (Fig. 2a; Fig. S5). Overall, we found that members of Ochrophyta, Ciliophora, Labyrinthulomycetes, and unicellular Opisthokonta were abundant across samples. Other commonly encountered taxa, which varied in abundance, included Fungi, Haptophyta, Chlorophyta, amoebozoa in the Thubulinea and Protosporangiida, Stramenophiles of the MAST-3 group, Apusomonadidae, and Cercozoa. Compared to the original oyster spat (T-1), a decrease in haptophytes was detected during the challenge experiment.

**Fig. 2.**
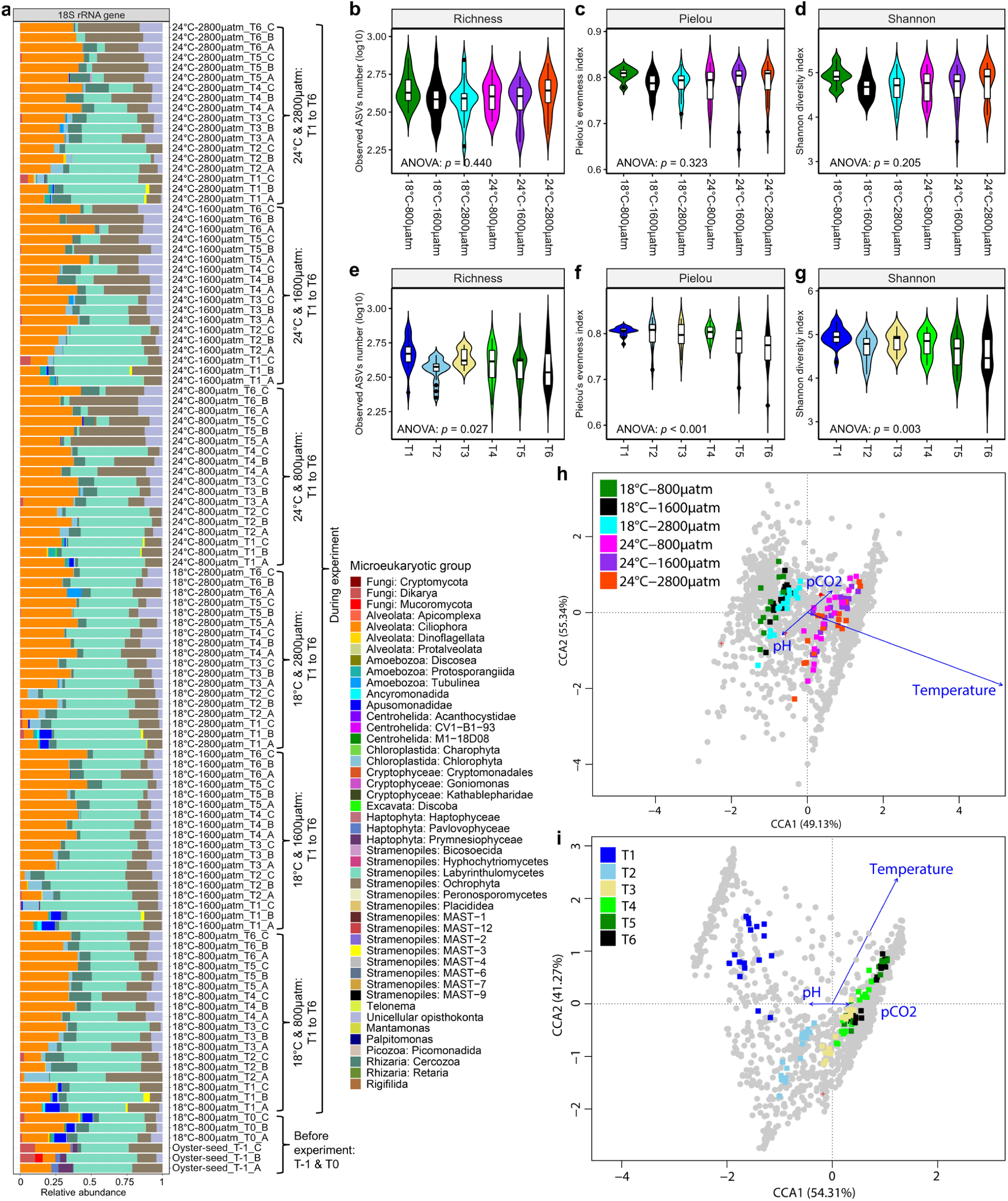
Variability of microeukaryotic community associated with Pacific oyster spat during the challenge experiment. **a** Relative abundance of microeukaryotic 18S rRNA gene sequences in oysters spat samples subjected to different temperatures (18℃, 24℃) and *p*CO_2_ (800, 1600, and 2800 µatm) conditions over the course of three weeks (T1 to T6). Additionally, oysters spat samples before the experiment (T-1: original oyster spat; T0: oyster spat sampled after 24-hour acclimation in tank-water before the experiment began) were also examined. **b-g** Alpha diversity (e.g., richness, Pielou’s *J*, and Shannon’s *H*) of microeukaryotic 18S rRNA genes across time and treatments. The violin plot is a smoothed density plot of the data, with the violin’s width representing the frequency of observations in that range of values. The line in the middle of the violin plot represents the data’s median, and the square indicates the data’s mean. **h & i** Canonical Correspondence Analysis (CCA) with measured abiotic variables within mesocosms (temperature, *p*CO_2_, and pH) as constrained variables to explain community changes of microeukaryotic 18S rRNA gene sequences associated with oysters between treatments (**h**) and time points (**i**). Grey points represent different microeukaryotic ASVs, and the square shapes indicate the oyster samples. Panel **b-d** shares the same color code as panel **h**, while panel **e-g** shares the same color code as panel **i**.

Estimates of microeukaryotic alpha diversity varied among treatments with richness ranging between 188 and 734 (average 412 ± 113 [mean ± SD, n = 108]; Fig. 2b,e), Pielou’s evenness index (*J’*) between 0.64 and 0.84 (average 0.79 ± 0.03 [mean ± SD, n = 108]; Fig. 2c,f), and the Shannon diversity index (*H’*) between 3.45 and 5.30 (average 4.73 ± 0.35 [mean ± SD, n = 108]; Fig. 2d,g). The differences were significant among time points for richness (ANOVA: *p* = 0.027), Pielou’s *J* (ANOVA: *p* < 0.001), and Shannon’s *H* (ANOVA: *p* = 0.003) (Fig. 2e-g), but not for temperature and *p*CO_2_ (Fig. 2b,d; Fig. S6).

The microeukaryotic community structure also varied across time and treatments (Fig. S4; Fig. S7-8; Fig. 2). Specifically, community composition differed significantly among the six temperature and *p*CO_2_ treatments (PERMANOVA: r^2^ = 0.093, *p* = 0.001), between the two temperatures (PERMANOVA: r^2^ = 0.023, *p* = 0.002), and across the six-time points (T1 to T6; PERMANOVA: r^2^ = 0.201, *p* = 0.001), but did not differ significantly among the three *p*CO_2_ levels (PERMANOVA: r^2^ = 0.018, *p* = 0.523). These significant differences are evident by PCoA and CCA, which showed that microeukaryotic communities from the same temperature (18℃, 24℃) or sampled at the same time point (T1 to T6) roughly clustered together (Fig. 2h-i; Fig. S8).

### Core constituents of the microbiome in oyster spat across time and treatments

For each combination of temperature and *p*CO_2_ treatment, the core microbiota (ASVs with a relative abundance >1% in >50% of samples) across all six time points comprised 13 prokaryotic and 38 microeukaryotic ASVs. The sum of the sequences assigned to these core taxonomic groups accounted for 0% to 50.2% (median = 17.3%) and 4.9% to 57.6% (median = 21.9%) of the relative abundance in prokaryotic and microeukaryotic communities, respectively (Fig. 3; Supplementary Results). Of these, there were seven and nine core prokaryotic ASVs (Fig. S9a-b), as well as 15 and 19 core microeukaryotic ASVs (Fig. S9c-d), at 18°C and 24°C, respectively. At 800, 1600, and 2800 *µ*atm *p*CO_2,_ there were seven, eight and seven core prokaryotic ASVs (Fig. S10a-b) as well as 13, 15, and 18 core microeukaryotic ASVs (Fig. S10c-d), respectively.

**Fig. 3.**
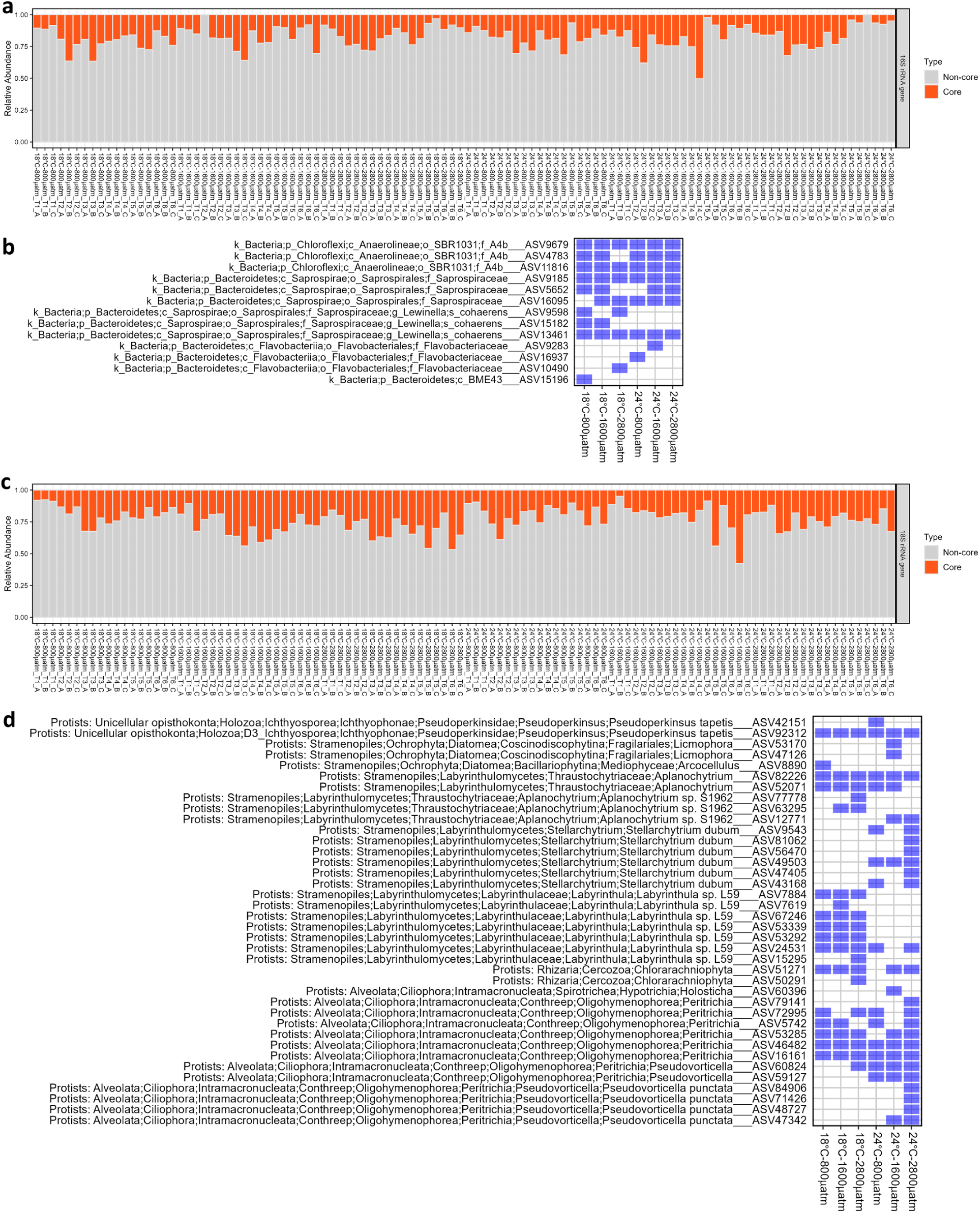
Core constituents of prokaryotic (a, b) and eukaryotic (c, d) microbiome in Pacific oyster spat samples across time in each treatment. **a & c** Relative abundance of core and non-core constituents of the prokaryotic (**a**) and eukaryotic (**c**) microbiota in oyster spat samples from the challenge study. **b & d** Presence/absence of these core constituents of the prokaryotic (**b**) and eukaryotic (**d**) microbiota in oyster spat samples of each treatment across time. Blue squares indicate the presence of the core taxa.

In addition, the core microbiota was examined at each time point for oyster samples across all six temperature and *p*CO_2_ treatments (Fig. S11-12). The results revealed that core taxa, consisting of 28 prokaryotic (Fig. S11) and 73 microeukaryotic (Fig. S12) ASVs, accounted for 0% to 59.2% (median = 27.2%) and 13.2% to 62.3% (median = 30.6%) of the relative abundance, respectively. The occurrence of core taxa varied through time (Fig. S11-12; Supplementary Results).

### Differentially occurring microbial taxa through time

LEfSe analysis revealed that different microbial ASVs occurred through time (Fig. 4a-b). For the prokaryotic microbiome (Fig. 4a), there was a clear shift in abundance from Bacteroidetes and Gammaproteobacteria in the genera *Fluviicola* and *Neptuniibacter*, respectively, in Week 1 (T1-T2), to Bacteroidetes in the genus *Lewinella* in Week 2 (T3-T4), and a resurgence of Gammaproteobacteria in the genus *Marinicella* in Week 3 (T5-T6), and subsequently by Betaproteobacteria in the genus *Methylotenera*. Within the microeukaryotes (Fig. 4b), a large portion of notable ASVs in Week 1 consisted of Labyrinthulomycetes in the genera *Stellarchytrium* and *Aplanochytrium*, Choanoflagellida in the genus *Acanthoeca*, and Chlorophyta in the genera *Nannochloropsis* and *Pycnococcus*. In Week 2, Cercozoa (genus *Ebria*) and diatoms (genus *Chaetoceros*) instead made up most of the differentially occurring taxa. A further shift towards differentially abundant ciliates (*Pseudovorticella* spp.), diatoms (*Licmophora* spp.), Ichthyosporea (*Pseudoperkinsus* spp.), and labyrinthulomycetes (*Labyrinthula* spp.) was observed in Week 3.

**Fig. 4.**
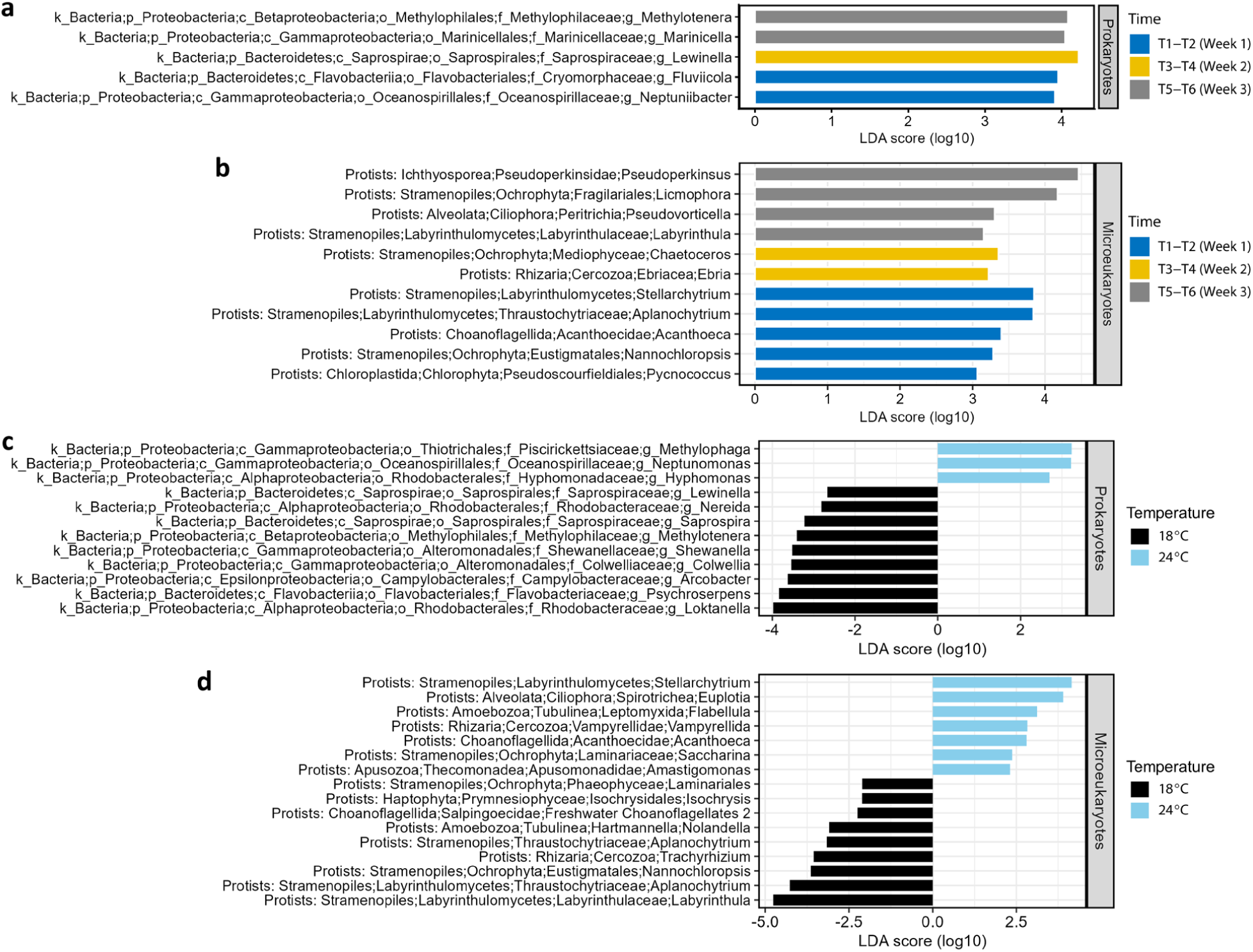
Differentially occurring microbial taxa in Pacific oyster spat samples. **a-d** Linear discriminant analysis Effect Size (LEfSe) reveals differentially (*p* < 0.05) detected prokaryotic (**a**, **c**) and microeukaryotic (**b**, **d**) genera in oyster spat samples across time (**a**, **b**) and between temperatures (**c**, **d**). Histograms show the LDA scores calculated for the difference in abundance among prokaryotic 16S (**a**, **c**) or microeukaryotic 18S (**b**, **d**) rRNA gene sequences binned to the genus level.

### Differentially occurring microbial taxa between temperatures

LEfSe analysis was carried out to determine which microorganisms may account for the effects of temperature on the prokaryotic and eukaryotic microbiomes of oyster spat (Fig. 4c-d). Genera within the Proteobacteria and Bacteroidetes were overrepresented at 18°C (Fig. 4c), including *Loktanella, Psychroserpens, Arcobacter, Colwellia, Shewanella, Methylotenera, Saprospira, Nereida,* and *Lewinella*. On the other hand, genera in the phylum Bacteroidetes, such as *Methylophaga, Hyphomonas,* and *Neptunomonas*, were the most differentially abundant taxa at 24°C. As well, several microbial eukaryotic taxa differed in abundance between temperatures (Fig. 4d), including the genera *Isochrysis*, *Laminariales*, *Nolandella*, *Aplanochytrium*, *Trachyrhizium*, *Nannochloropsis*, *Aplanochytrium*, and *Labyrinthula* at 18°C, while *Stellarchytrium*, *Euplotia*, *Flabellula*, *Vanpyrellida*, *Acanthoeca*, *Saccharina*, and *Amastigomonas* were overrepresented at 24°C. No differentially abundant taxa were found among the three *p*CO_2_ conditions.

### Occurrence of potentially pathogenic microbial species across time and treatments

We screened the prokaryotic and microeukaryotic data for taxa that include potential pathogens of shellfish, including oysters (see pathogen lists in Bower (2010)). In *M. gigas* spat, rRNA gene sequences of the putative oyster pathogen, *Uronema marinum* (Plunket and Hidu 1978), were abundant at T-1, T0, and T1, but decreased to mostly undetectable levels at later time points regardless of temperature and *p*CO_2_ treatment (Fig. 5; Supplementary Results). Additionally, the relative abundance of 18S rRNA gene sequences from *Pseudoperkinsus tapetis*, a well-known clam pathogen (Villalba et al. 2004), increased with time points in all six treatments (Fig. 5; Supplementary Results).

**Fig. 5.**
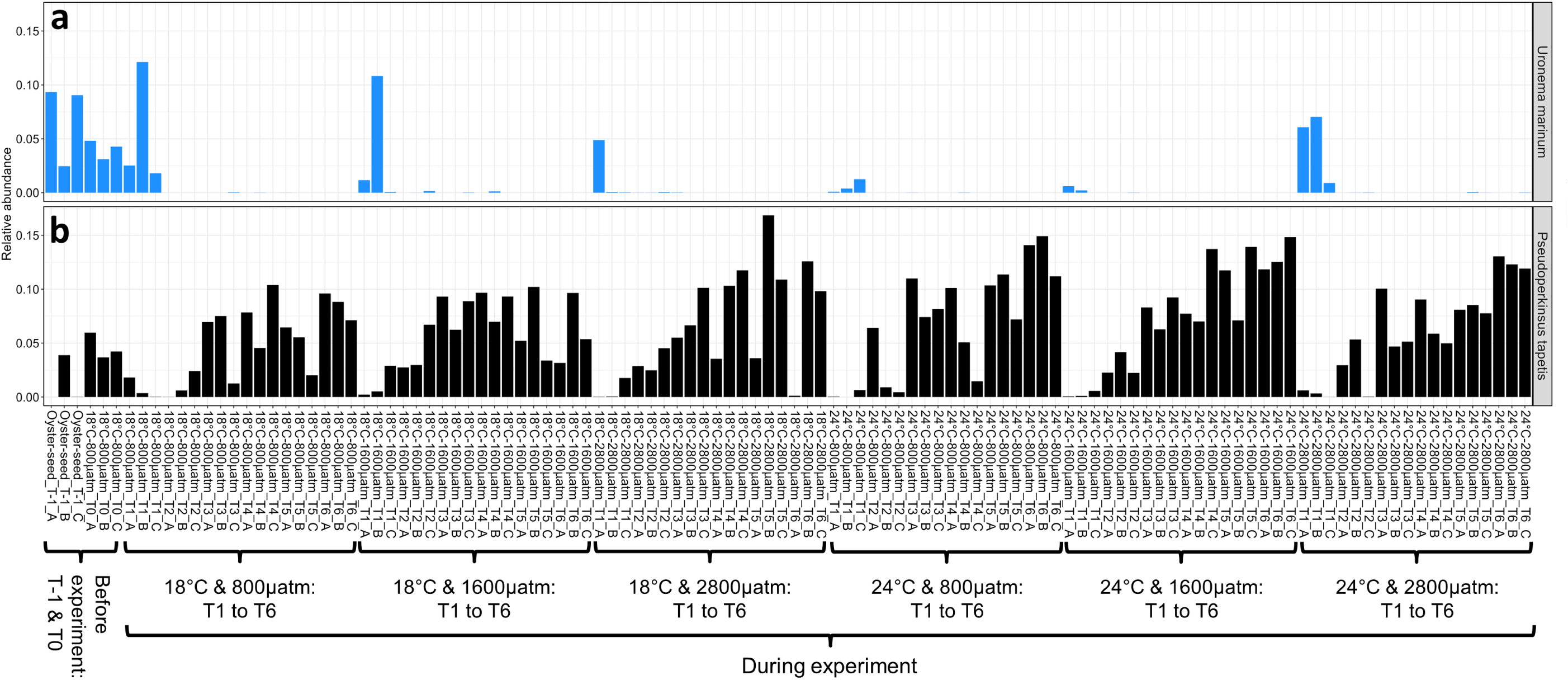
Relative abundances of potentially pathogenic microbes in Pacific oyster spat. **a-b** Relative abundances of the taxa *Uronema marinum* (**a**) and *Pseudoperkinsus tapetis* (**b**) in oyster spat samples across time under different temperature and *p*CO_2_ conditions. Additionally, oyster spat samples prior to the start of the experiment (T-1: original oyster seed; T0: oyster spat sampled after 24-hour acclimation in tank-water) were also examined.

### Pairwise correlations between microbial communities and the measured abiotic and biotic variables in Pacific oyster spat samples

Over the course of six time points, our Pearson correlation analysis revealed a range of pairwise correlation coefficient (*r*), from -0.976 (between *p*CO_2_ and pH) and 0.821 (between 18S_Pielou and 18S_Shannon) (Fig. 6), among the measured abiotic (temperature, *p*CO_2_, pH, and time [days since the experiment started]) and biotic (prokaryotic and microeukaryotic alpha-diversity indices, including the observed ASV richness, Pielou’s evenness index and Shannon’s diversity index) variables of the oyster spat samples (n = 108). The Mantel test revealed significant (*p* < 0.05) correlations between the prokaryotic community composition and all measured variables, with the exception of *p*CO_2_, pH and the richness of microeukaryotes (Fig. 6). The microeukaryotic community composition was significantly (*p* < 0.01) correlated with temperature, time and microeukaryotic alpha-diveristy indices including richness, Pielou’s evenness index and Shannon’s diversity index (Fig. 6).

**Fig. 6.**
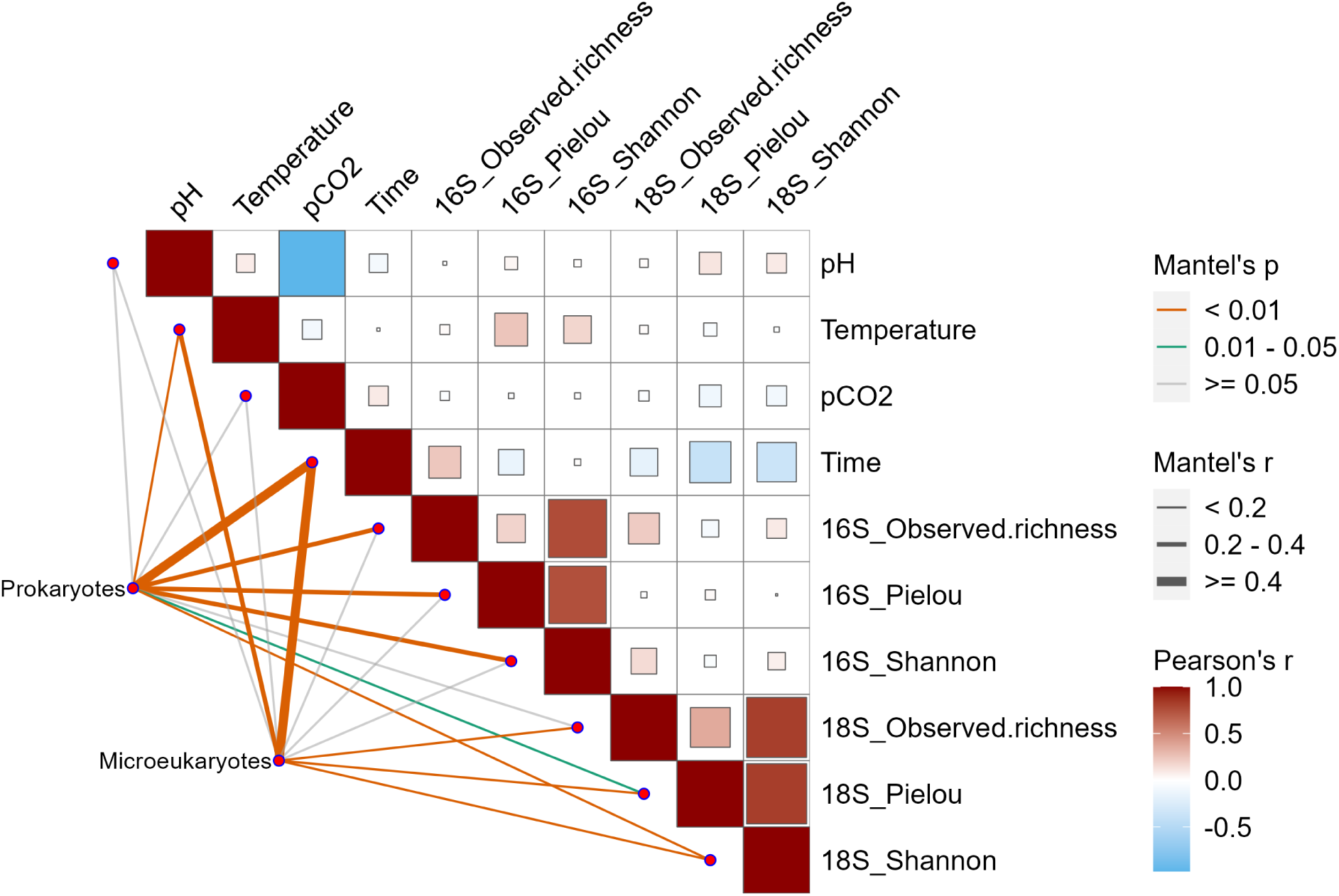
Pairwise correlations between microbial communities and the corresponding abiotic and biotic variables in Pacific oyster spat samples. These relationships are evaluated using Mantel test, with Pearson correlation analysis implemented on the pairwise Euclidean distances computed between abiotic (temperature, *p*CO_2_, pH, and time [days since the experiment started]) and biotic (the alpha-diversity indices of prokaryotic and microeukaryotic communities, respectively, including the observed ASV richness, Pielou’s evenness index and Shannon’s diversity index) variables of the oyster spat samples (n = 108). Line width denotes the magnitude of Mantel’s correlation coefficient, while color indicates the statistical significance of the correlation. Pearson correlation coefficients of the Euclidean distances between measured biotic and abiotic variables were shown in the heatmap. The size of each square in the heatmap corresponds to the absolute Pearson correlation coefficient value, with the color gradient indicating the correlation value, ranging from red (high correlation) to blue (low correlation).

## Discussion

Documenting the effects on the oyster microbiome of changes in ocean temperature and acidity is critical to understanding how the productivity, health, and survival of wild and farmed oysters may be affected by future climate scenarios. This includes the eukaryotic microbiome, which is rarely examined (Bharti and Grimm 2021; Paillard et al. 2022), but comprises a diverse and significant proportion of host-associated microbes (Laforest-Lapointe and Arrieta 2018; Mann et al. 2020). As such, documenting changes in the prokaryotic and eukaryotic microbiome of oysters can reveal holistic effects of environmental change on the microbiota. In this study, we examined the relationship between environmental factors linked to climate change and the microbiome associated with Pacific oyster spat. Our results showed that the prokaryotic and eukaryotic microbiome of oyster spat is influenced by temperature and maturation, but not by *p*CO_2_. These findings have important implications for understanding how the oyster microbiome and hence populations may be affected by changes in ocean warming and acidification, and demonstrate the resilience of the oyster-spat microbiome to changing environmental conditions.

### Documenting the prokaryotic and eukaryotic microbiome of M. gigas spat

Oysters are filter feeders (Dupuy et al. 1999) that consume phytoplankton (autotrophic microbes), heterotrophic protists, zooplankton (e.g., copepods), planktonic eggs and larvae, bits of detrital matter, and likely, bacteria (Dupuy et al. 1999; Kamiyama 2011; Weissberger and Glibert 2021). This filter-feeding strategy was especially evident in the eukaryotic microbiome, in which 5.8% to 51.2% of 18S rRNA gene sequences belonged to microalgae, including diatoms, chlorophytes, cryptophytes, and haptophytes (Fig. 2; Fig. S5).

Another dietary component, heterotrophic protists, graze microbes and autotrophic picoplankton that are too small to be effectively retained by oysters (Barillé et al. 1993; Riisgård 1988; Shumway et al. 1985). Hence, heterotrophic protists are a trophic link between picoplankton and benthic filter-feeders (Dupuy et al. 1999; Kamiyama 2011; Le Gall et al. 1997). Our results show that heterotrophic protists belonging to the Ciliophora, Labyrthulomycetes, Cercozoa, and unicellular Opisthokonta were abundant in oyster spat (Fig. 2; Fig. S5), but it is unclear which protists are food and which are the intrinsic residents in oysters.

Ciliates are protozoa that feed on bacteria and small eukaryotes and which in turn serve as food for oysters (Le Gall et al. 1997), providing both nutrients and carbon (Dupuy et al. 1999; Kamiyama 2011). In our experiments, most ciliates belonged to the ciliophoran subclass *Peritrichia* (e.g., genus *Pseudovorticella*; Fig. S5), consistent with a study of adult Pacific oysters in France that found peritrichous ciliates (genus *Trichodina*) were abundant (Clerissi et al. 2020). Notably, some ciliates can cause disease in oyster spat, such as those in the subclass *Scuticociliatia* (e.g., *Miamiensis avidus*, *Paranophrys* sp., *Uronema* sp.), which were present in oyster spat samples from a mortality event in in British Columbia, Canada (Zhong et al. 2021).

Labyrinthulomycetes are fungal-like protists that are widely present in the marine environment, particularly in metazoans and metaphytes (Collier et al. 2017). Most labyrinthulomycetes are saprotrophic feeders associated with detritus such as fallen mangrove leaves, decomposing algae, and fecal pellets of marine invertebrates (Pan 2016; Raghukumar 2002; Tsui et al. 2009). Although labyrinthulomycetes were reported to cause diseases in animals (e.g., quahog parasite X disease in hard-shell clams caused by *Thraustochytrid* sp.; Ragan et al. 2000) and seagrass (e.g., wasting disease caused by *Labyrinthula* sp.; Bigelow et al. 2005; Muehlstein et al. 1991), labyrinthulomycetes are often seen to live as commensals or mutualists within the guts and tissues of healthy marine invertebrates (Pan 2016). In Pacific oyster spat, most labyrinthulomycetes detected belonged to the genera *Aplauochitrim*, *Labyrinthula* and *Stellachytrium* (Fig. S5), although there was no indication that they were associated with disease in our study, suggesting they may be involved in the decomposition of organic matter and contribute to the nutrition of the oyster spat.

Across all treatments and time points, the prokaryotic microbiome mainly consisted of members of the phyla Bacteroidetes and Proteobacteria, accompanied by a sizable proportion of Chloroflexi and Planctomycetes (Fig. 1; Fig. S1). These results are consistent with research that discovered an abundance of Bacteroidetes and/or Proteobacteria in *M. gigas* adults and spat (Fernandez-Piquer et al. 2012; Li et al. 2017). Although the role of bacteria in these phyla is unclear in oysters, their dominance of the prokaryotic community could be a result of a symbiotic relationship, in which proteobacteria aid in the digestion of cellulose and agar consumed by *M. gigas* as well as provide fixed nitrogen to their host (Li et al. 2017; Trabal Fernández et al. 2014; Zehr et al. 2003). Bacteroidetes are also known to be important commensals of the gut due to their ability to break down plant material (Li et al. 2017; Thomas et al. 2011; Trabal Fernández et al. 2014).

### The core microbiome of M. gigas spat is comprised of a small number of relatively abundant taxa

The taxa comprising the core microbiome in oyster spat provides insights into the potential role of the microbiome in the adaptation of oyster spat to a changing environment (Risely 2020; Shade and Handelsman 2012). Here, we found that the core microbiome of spat comprised only a few taxa that represented, at most, 0.2% of the total number of ASVs, but on average accounted for 23.5% to 31.1% of the total relative abundance of prokaryotes and eukaryotes, respectively. This suggests a small subset of the microbial assemblage of oyster spat may be important for maintaining the functional stability of the microbiome across time (Fig. S11-12) and treatments (Fig. 3; Fig. S9-10). The core taxa (ASVs) comprised Chloroflexi, Bacteroidetes, Ochrophyta, Labyrinthulomycetes, Cercozoa, Ciliophora, Apusomonadidae, Chlorophyta, Amoebozoa, and Ichthyosporea, with their occurrence and relative abundances varying over time and among treatments (Fig. 3; Fig. S9-12). The variability in some core taxa across different temperatures and *p*CO_2_ treatments suggests that although defined as core, they were sensitive to different environmental conditions (Fig. 3; Fig. S9-10). Moreover, some members of the core microbiome changed over time, with different taxa becoming more or less abundant (Fig. S11-12), emphasizing that not all core taxa were stable as the oysters aged. Interestingly, taxa in the Chloroflexi family A4b (e.g., ASV9679, ASV11816, ASV9185), Bacteroidetes family *Saprospiraceae* (e.g., ASV9185, ASV13461), Labyrinthulomycetes genera *Labyrinthula* (e.g., ASV24531) and *Aplanochytrium* (e.g., ASV82226), and the Ichthyosporea species, *Pseudoperkinsus tapetis* (e.g., ASV92312), were core constituents (i.e. ASVs with a relative abundance ≥1% in ≥50% of samples for a treatment or time-point) across all six combinations of temperature and *p*CO_2_ treatments and time points (Fig. 3; Fig. S9-12). This finding implies that, regardless of environmental conditions or age, these taxa are likely key components in *M. gigas* spat.

Despite the importance of characterizing the core microbiome as a means of discovering microorganisms that hold important relationships with their host (D Ainsworth et al. 2015; King et al. 2019c), the core prokaryotic microbiome of oysters has only been reported in a few studies (King et al. 2019a; King et al. 2019c; Stevick et al. 2021; Unzueta-Martínez et al. 2022). For example, eight OTUs were unique to healthy Australian Pacific oysters, and included the taxa *Arcobacter* and *Vibrio fortis* (King et al. 2019a). In contrast, we found that a large majority of the core ASVs were assigned to the Bacteroidetes, though their functional importance in *M. gigas* spat remains unknown. We also report on the eukaryotic microbiome of *M. gigas* spat. Further investigations are needed to determine if the microbes making up the core prokaryotic and microeukaryotic microbiomes found in this study are also representative of the microbiomes in other Pacific oysters worldwide, and their persistence under different environmental conditions.

### The microbiome of M. gigas spat changes over time

Early oyster spat undergo major physiological changes as they transition to a benthic lifestyle (King et al. 2019a). Accordingly, we hypothesized that the microbiome of spat would shift over time as it matured along with its host, as shown previously (Cho et al. 2023; Cho 2019; Trabal Fernández et al. 2014). Across both the prokaryotic and eukaryotic microbiomes, there were significant compositional changes during the three weeks of sampling (PERMANOVA: prokaryotes, r^2^ = 0.199, *p* = 0.001; microeukaryotes, r^2^ = 0.201, *p* = 0.001; Mantel test: prokaryotes, r = 0.567, *p* = 0.001; microeukaryotes, r = 0.558, *p* = 0.001) (Table S3; Fig. 6; Fig. 1h; Fig. 2h; Fig. S4; Fig. S8). Such significant changes were likely driven by both richness and evenness (Fig. 1e-g; Fig. 2e-g). These changes were evident through LEfSe analysis, which identified a variety of differentially occurring microbial taxa week by week (Fig. 4a-b). Various prokaryotic and microeukaryotic taxa fared better in the younger spat, such as *Fluviicola*, *Neptuniibacter*, *Stellarchytrium*, *Aplanochytrim*, *Acanthoeca*, *Nannochloropsis*, and *Pycnococcus*. In comparison, others thrived in the older spat, including *Methylotenera, Marinicella, Pseudoperkinsus, Pseudovorticella, Labyrinthula,* and *Licmophora*.

Further evidence that the microbiome of spat changes as the host develops is that the potential pathogen, *Uronema marinum* (Plunket and Hidu 1978), was abundant at T-1, T0 and T1, and accounted for up to 12.1% of the relative abundance of ribosomal rRNA gene sequences, but was mostly undetectable at later time points, regardless of treatment (Fig. 5). Although the reasons for change in the overall microbiome is unknown, it is clearly associated with a decline in the relative abundance of *U. marinum*, suggesting that the maturation of the microbiome aids in pathogen defense, potentially through competition for nutrients, maintenance of immune homeostasis, and antimicrobial activity (Desriac et al. 2014; Dupont et al. 2020; King et al. 2019b). Evidence that spat age drives shifts in microbial composition can be found in data from experiments in which spat were reared under constant environmental conditions; yet, age was the best predictor of microbial composition (Cho 2019; Cho et al. 2023).

In addition, we detected a Perkinsus-like organism, *Pseudoperkinsus tapetis*, that was abundant in most Pacific oyster spat samples, and accounted for up to 16.9% of the 18S sequences, although varying across time and treatments (Fig. 5). Moreover, *Pseudoperkinsus tapetis*, belongs to the unicellular opisthokont lineage *Ichthyosporea*, and occurs in the guts of marine invertebrates ranging from oysters to sea cucumbers (Clerissi et al. 2020; Marshall and Berbee 2010); it has been shown to cause perkinsosis disease in clams (Villalba et al. 2004). Another Perkinsus-like organism, *Perkinsus marinus,* causes devastating mortality in eastern oysters (*C. virginica*) (Andrews 1988; Cook et al. 1998; Villalba et al. 2004). However, we did not detect mortality of oyster spat in this experiment (J. F. Finke unpubl.), and thus *P. tapetis* may not cause disease in oyster spat, but rather be commensal or a mutualist.

### Warming seawater drives microbiome shifts in M. gigas spat

Temperature significantly influenced the composition of the prokaryotic and eukaryotic microbiome over the three-week challenge experiment (PERMANOVA: prokaryotes, r^2^ = 0.022, *p* = 0.002; microeukaryotes, r^2^ = 0.023, *p* = 0.002; Mantel test: prokaryotes, r = 0.073, *p* = 0.001; microeukaryotes, r = 0.242, *p* = 0.001) (Table S3; Fig. 6; Fig. 1i; Fig. 2i; Fig. S4; Fig. S8), although the r^2^ value in PERMANOVA is relatively low and the temperature effect on alpha-diversity was not significant (Fig. S2d-f; Fig. S5d-f). Various taxa were specific to each treatment (Fig. 4c-d), which could be because of different temperature optima among taxa. For example, prokaryote genera that were more abundant at 24°C, included *Methylotenera* (Kalyuzhnaya et al. 2006), *Neptunomonas* (Diéguez et al. 2017) and *Methylophaga* (Villeneuve et al. 2013), genera that comprised some species with optimum growth temperatures ranging from 23 to 30°C (Lyubetsky et al. 2020). In contrast, prokaryotic genera that were abundant at the 18°C treatment included *Arcobacter* (Van Driessche and Houf 2008), *Colwellia* (Techtmann et al. 2016), and *Saprospira* (Saw et al. 2012); these genera contain members that grow well at 18°C, and can tolerate temperatures as low as 4°C. Other studies have found that temperature and heat stress can significantly alter the prokaryotic microbiome of Pacific oysters (Green et al. 2019; Lokmer and Mathias Wegner 2015); here, we show that the eukaryotic microbiome is also affected. Consequently, our results highlight the importance of considering microbial eukaryotes when assessing the potential impacts of microbiome changes on Pacific oysters, especially in the context of rising sea temperatures.

Given that temperature also affects the prokaryotic microbiome of other oyster species, such as *Saccostrea glomerata* (Scanes et al. 2021a; Scanes et al. 2021b; Scanes et al. 2021c), it seems evident that ocean warming will affect oyster microbiomes, in general, with potential consequent effects on oyster health. The implications of temperature-induced changes in the microbiome for long-term oyster health and resilience of populations remain to be elucidated.

### Resilience of M. gigas spat microbiome to acidification

We grew Pacific oyster spat at *p*CO_2_ levels of 800, 1600, and 2800 *μ*atm (e.g., resulting in pH mean ± SD in tank water at the 18°C treatment: 7.79 ± 0.09 [n = 30], 7.50 ± 0.04 [n = 30] and 7.24 ± 0.04 [n = 30], respectively; further details are in Table S1) to assess the effect of acidification on the spat microbiome. Previous studies have shown that changes in pH as a result of *p*CO_2_, influence the physiology of marine organisms such as corals and oysters (Alma et al. 2020; van Hooidonk et al. 2014; Waldbusser et al. 2015), and also play a role in shifting the composition of the microbiome (Scanes et al. 2021a; Scanes et al. 2021b; Scanes et al. 2021c; Unzueta-Martínez et al. 2021). Therefore, we expected *p*CO_2_ to affect the biology of the spat, and thus influence the microbiome. However, in our study, *p*CO_2_ level was not significantly related to changes in alpha-diversity or composition of either the prokaryotic or eukaryotic microbiomes (PERMANOVA: prokaryotes, r^2^ = 0.019, *p* = 0.386; microeukaryotes, r^2^ = 0.018, *p* = 0.523; Mantel test: *p* > 0.05 for both prokaryotes and microeukaryotes) (Table S3; Fig. 6; Fig. S2a-c; Fig. S6a-c), even though a *p*CO_2_ level of 2800 *μ*atm equates to a pH drop of ∼0.55 (e.g., the 18°C treatment), which is greater than the predicted pH drop of 0.29 in the ocean by the year 2100 (Bindoff et al. 2019). In contrast to our observations, challenge experiments with adult eastern oysters (*Crassostrea virginica*, pH drop ∼0.7; Unzueta-Martínez et al. 2021) and Sydney rock oysters (*Saccostrea glomerata*, pH drop ∼0.34; Scanes et al. 2021a; Scanes et al. 2021b; Scanes et al. 2021c) showed that *p*CO_2_-mediated acidification affected the microbiome. Possibly, the lack of an observable effect of acidification on the spat microbiome could be attributed to resilience to changes in *p*CO_2_ and pH, as has been reported in terms of survival across multiple oyster species (Ginger et al. 2013; Lawlor and Arellano 2020). Another possibility is that because the microbiome of oyster spat is less established, it is also less specialized, and hence, more resilient to changes in acidity compared to adults. Although there can be transgenerational effects that increase the resilience of oysters to acidification (Lim et al. 2021b; Scanes et al. 2023), this was not a factor in our study, as the broodstock was not exposed to high CO_2_ conditions. It is possible that longer exposure of the spat may have resulted in a *p*CO_2_ effect on the microbiome, but intuitively the effects of changes in acidity would be expected to be manifested fairly quickly; moreover, we are unaware of studies showing that changes in acidification do not affect the composition of the oyster microbiome on the short-term, but affect the microbiome on longer time scales. Ultimately, the resilience of the spat microbiome to changes in ocean acidity likely contributes to the maintenance of oyster health under varying environmental conditions.

## Conclusion

Increased atmospheric carbon dioxide is causing acidification and warming of the oceans, which is associated with episodic changes in temperature and *p*CO_2_ at local scales that can directly affect wild and cultured oyster populations. In order to assess the potential impacts of these environmental changes on early-development oysters, we subjected Pacific oyster (*M. gigas*) spat to different combinations of temperature and *p*CO_2_ over a period of three weeks, aiming to evaluate how these conditions influence the developing oyster microbiome. The results showed that warmer temperature had a significant influence on the oyster microbiome, while acidification did not.

Regardless of temperature or *p*CO_2_, the prokaryotic and eukaryotic microbiome changed across time, with potentially pathogenic ciliates (*Uronema marinum*) greatly reduced in all treatments. Moreover, we show that temperature affects both the prokaryotic and eukaryotic microbiome of oyster spat, emphasizing that microbial eukaryotes also need to be considered when contemplating how changes in the microbiome may affect oyster health. This work highlights the short-term impacts that changes in *p*CO_2_ and temperature can have on the microbiome in developing oyster spat.

## Data availability statement

All next-generation sequencing data generated in this study have been deposited in the NCBI Sequence Read Archive (SRA) under the accession numbers SRR22282694 to SRR22282807 and SRR22282906 to SRR22283019 for 16S and 18S rRNA gene sequences, respectively. The authors declare that all other data supporting the findings of this study are available within the paper and/or the associated supplementary files.

## Acknowledgments

We thank Claire Rycroft from Pacific Biological Station (Fisheries and Oceans Canada), as well as Iria Gimenez, Josianne Haag, Jacob Etzkorn, and Jonathan Bergshoeff from the Hakai Institute for assisting in the challenge experiments. Walcan Seafoods Ltd. provided access to their pumping system for seawater to initiate the experiment. This project was funded by the Gordon and Betty Moore Foundation, GBMF#5600, to C.A.S., K.M.M. and S.P.O. Support for the challenge experiments was provided by the Tula Foundation and the Marnalab facilities at the Hakai Institute Quadra Island Ecological Observatory. Additional support was provided by an infrastructure award from the Canada Foundation for Innovation and British Columbia Knowledge Development Fund (Project #25412 to C.A.S.).

## Author Contribution Statement

K.X.Z. and M.D. wrote the draft manuscript, which was revised by K.X.Z and C.A.S. following input from all co-authors. The experiments were conceived and designed by C.A.S., A.M.C., K.M.M. and S.P.O. with input from R.S., C.D.G.H., J.F.F., B.J.G.S., A.V.H., T.J.G. and K.X.Z. The challenge experiments were set up by B.C., K.R., M.F., J.F.F., C.A.S., A.M.C. and A.V.H.; oyster spat was provided by T.J.G. Operation of the experimental system and sample collection was done by B.C., K.R. and M.F. K.R. and B.C. did the *p*CO_2_ measurements. K.X.Z. processed the samples, prepared the sequencing libraries, conducted the bioinformatic and statistical analyses, and the interpretation. Except for the corresponding authors, all co-authors are listed alphabetically according to family name.

## Supplementary Information

### 1. Supplementary Methods

#### 1.1. Amplicon-sequencing library preparation for 16S rRNA genes

Deep sequencing of 16S rRNA gene amplicons was employed to profile prokaryotic microbiota associated with oysters. The preparation of 16S amplicon libraries was adapted from the online Illumina protocol (Amplicon et al. 2013) with several modifications. Briefly, two successive runs of PCR were performed. The first PCR generated amplicons of the ∼292-bp 16S rRNA gene between the V4 region using the modified primers 515F-Nxt and 806R-Nxt (Table S2). Compared to 515F and 806R (Apprill et al. 2015), these modified primers incorporate overhanging adapter sequences (Table S2) that are compatible with Illumina index and sequencing adapters. These modifications enabled the use of Illumina Nextera XT indexes as forward and reverse primers to create the dual-indexed amplicon libraries in the second PCR.

In the first PCR, the 25-μL reaction mix was made with 1X PCR buffer, 4 mM MgCl_2_, 50 µg of Bovine Serum Albumin (Invitrogen), 200 mM of each dNTP (Invitrogen), 0.4 µM of each primer, 0.5 U of Q5® high fidelity polymerase (NEB) and approximately 10 ng of genomic DNA template. The program started with an initial denaturation of three minutes at 95°C, followed by 29 cycles of denaturation at 95°C for 45 s, annealing at 50°C for 60 s, elongation at 72°C for 90 s, and a final elongation for 10 minutes at 72°C. PCR products were purified using SPRIselect beads (Beckman Coulter) with a ratio of 1:1 for beads:product to remove fragments less than 200bp (e.g., primer dimers).

These purified amplicons created by the first PCR were then used as the template for the index-PCR (i.e., second PCR) to generate an amplicon-sequencing library. The parameters for the index-PCR were the same as the online Illumina protocol (Amplicon et al. 2013) but used the Q5® high fidelity polymerase (NEB) instead. The index-PCR started with an initial denaturation of three minutes at 95°C, followed by 12 cycles of denaturation at 95°C for 30 s, annealing at 55°C for 30 s, elongation at 72°C for 30 s, and a final elongation for 10 minutes at 72°C. The 25-μL reaction mix was made with 1X PCR buffer, 4 mM MgCl_2_, 200 mM of each dNTP (Invitrogen), 2.5 µL of each index primer (N7XX and S5XX of Nextera® XT Index Kit), 1 U of Q5® high fidelity polymerase (NEB) and 2.5 µL of purified amplicons from the first PCR.

The amplicon-sequencing library was obtained by purifying the index-PCR product using SPRIselect beads (Beckman Coulter) with a bead:product ratio of 1:1. The amplicon-sequencing library was quantified using a Qubit® dsDNA HS Assay Kit (Invitrogen), and the fragment size was determined using Agilent High Sensitivity DNA Kit (Agilent) on an Agilent 2100 Bioanalyzer System. Equimolar amounts of these barcoded and purified amplicon-sequencing libraries were pooled and sent to The University of British Columbia’s BRC-Seq Next-Gen Sequencing Core for sequencing using the MiSeq platform with 300-bp pair-end chemistry (Illumina).

### 2. Supplementary Results

#### 2.1. Core constituents of microbiome in oyster spat across time and treatments

ASVs with a relative abundance >1% and a >50% occurrence rate across examined oyster samples (e.g., by treatments or time points) were categorized as core constituents of the oyster spat microbiota (Miller et al. 2020; Unzueta-Martínez et al. 2022). For each combination of temperature and *p*CO_2_ treatment, the core microbiota for oyster samples across all six-time points was examined (Fig. 3). The results showed that 13 prokaryotic and 38 microeukaryotic ASVs are core constituents, which account for up to 50.2% and 57.6% of the relative abundance, respectively (Fig. 3). These constituents belong to the Chloroflexi family A4b, Bacteroidetes families *Saprospiraceae* and *Flavobacteriaceae* and class BME43, Ochrophyta genera *Licmophora* and *Arcocellulus*, Labyrinthulomycetes genera *Labyrinthula* and *Stellarchytrium*, Cercozoa class *Chlorarachniophyta*, Ciliophora subclass *Peritrichia*, and Ichthyosporea species *Pseudoperkinsus tapetis*. The occurrence of these core taxa varied among treatments (Fig. 3). Of these, seven and nine core prokaryotic ASVs (Fig. S9a-b) as well as 15 and 19 core microeukaryotic ASVs (Fig. S9c-d) were detected in two temperature conditions, 18°C and 24°C, respectively. Seven, eight and seven core prokaryotic ASVs (Fig. S10a-b) as well as 13, 15, and 18 core microeukaryotic ASVs (Fig. S10c-d) were detected in three *p*CO_2_ conditions, 800, 1600, and 2800 *µ*atm, respectively.

In addition, the core microbiota was examined at each time point for oyster samples across all six temperature and *p*CO_2_ treatments (Fig. S11-12). The results revealed that core taxa, consisting of 28 prokaryotic (Fig. S11) and 73 microeukaryotic (Fig. S12) ASVs, can make up to 59.2% and 62.3% of the relative abundance, respectively. These taxa belong to the Proteobacteria (particularly species *Neptuniibacter caesariensis*), Planctomycetes of order CL500-15, Chloroflexi family A4b, Bacteroidetes families *Saprospiraceae*, *Rhodothermaceae* and *Flavobacteriaceae* and class BME43, Ochrophyta genera *Nannochloropsis*, *Licmophora* and *Chaetoceros*, Labyrinthulomycetes of genera *Aplanochytrium*, *Labyrinthula* and *Stellarchytrium*, Cercozoa class *Chlorarachniophyta*, Ciliophora subclass *Hypotrichia*, *Euplotia* and *Peritrichia*, Ichthyosporea species *Pseudoperkinsus tapetis*, Apusomonadidae species *Amastigomonas* sp. 2 Bamfield, Chlorophyta species *Pycnococcus provasolii*, and Amoebozoa genus *Protosporangium*. The occurrence of these core taxa varied among time points (Fig. S11-12).

### 2.2. Occurrence of potentially pathogenic microbial species across time and treatments

We screened the prokaryotic and mciroeukaryotic taxa for the known putative pathogenic species of shellfish (see pathogen lists in Bower (2010)), including oysters. In *M. gigas* spat, we detected the putative oyster pathogens, *Uronema marinum* (Plunket and Hidu 1978). They occurred abundantly in samples at T-1, T0, and T1, but decreased to mostly undetectable levels at later time points regardless of temperature and *p*CO_2_ treatments (Fig. 5). The proportion of 18S rRNA gene sequences assigned to *U. marinum* is 6.949 ± 3.888% (mean ± SD, n = 3) in original oyster spat (T-1), 4.070 ± 0.867% (mean ± SD, n = 3) at T0, 2.783 ± 3.827% (mean ± SD, n = 18) at T1, 0.016 ± 0.038% (mean ± SD, n = 18) at T2, 0.006 ± 0.012% (mean ± SD, n = 18) at T3, 0.009 ± 0.030% (mean ± SD, n = 18) at T4, 0.004 ± 0.015% (mean ± SD, n = 18) at T5, and 0.001 ± 0.005% (mean ± SD, n = 18) at T6.

Furthermore, we detected *Pseudoperkinsus tapetis* in *M. gigas* spat, which is a Perkinsus-like unicellular opisthokonts that has been shown to cause perkinsosis disease in clams (Villalba et al. 2004). The proportion of 18S rRNA gene reads associated with *P. tapetis* is 1.301 ± 2.239% (mean ± SD, n = 3) in original oyster spat (T-1), 4.619 ± 1.199% (mean ± SD, n = 3) at T0, 0.556 ± 0.800% (mean ± SD, n = 18) at T1, 2.778 ± 2.077% (mean ± SD, n = 18) at T2, 7.371 ± 2.366% (mean ± SD, n = 18) at T3, 7.741±3.124% (mean ± SD, n = 18) at T4, 8.347 ± 3.836% (mean ± SD, n = 18) at T5, and 10.162 ± 4.028% (mean ± SD, n = 18) at T6, with the relative abundance varied across time points and treatments (Fig. 5).

## 4. Supplementary tables and figures

### 4.1. Supplementary tables list

- Table S1: Summary of mean experimental conditions and standard deviations in mesocosm chambers determined from discrete seawater samples collected across six sampling time points during the challenge experiment.
- Table S2: List of primers used in this study.
- Table S3: Dissimilarity of Pacific oyster spat associated prokaryotic and microeukaryotic community composition between two temperatures (18℃, 24℃) or among three *p*CO_2_ conditions (800, 1600, and 2800 µatm) across time revealed using permutational multivariate ANOVA (PERMANOVA) in the challenge experiment.

### 4.2. Supplementary figures list

- Fig. S1: Genus-level relative abundance of prokaryotes in Pacific oyster spat samples across time and between different temperature and *p*CO_2_ conditions.
- Fig. S2: Alpha diversity of prokaryotic 16S rRNA genes across temperature and *p*CO_2_ treatments in Pacific oyster spat.
- Fig. S3: Pearson correlation coefficients between prokaryotic communities of Pacific oyster spat across time and treatments.
- Fig. S4: Principal Coordinate Analyses (PCoA) of the community structure of the prokaryotic 16S rRNA gene sequences in oyster spat samples subjected to different temperature and pCO2 treatments across time.
- Fig. S5: Genus-level relative abundance of microeukaryotes in Pacific oyster spat samples across time and between different temperature and *p*CO_2_ conditions.
- Fig. S6: Alpha diversity of microeukaryotic 18S rRNA genes across temperature and *p*CO_2_ treatments in Pacific oyster spat.
- Fig. S7: Pearson correlation coefficients between microeukaryotic communities of Pacific oyster spat across time and treatments.
- Fig. S8: Principal Coordinate Analyses (PCoA) of the community structure of the microeukaryotic 18S rRNA gene sequences in oyster spat samples subjected to different temperature and pCO2 treatments across time.
- Fig. S9: Core constituents of prokaryotic and eukaryotic microbiota in Pacific oyster spat samples across time in each temperature treatment.
- Fig. S10: Core constituents of prokaryotic and eukaryotic microbiota in Pacific oyster spat samples across time in each *p*CO_2_ treatment.
- Fig. S11: Core constituents of the Pacific oyster spat prokaryotic microbiota among different treatments at each time point.
- Fig. S12: Core constituents of the Pacific oyster spat eukaryotic microbiota among different treatments at each time point.

**Table S1.** Summary of mean experimental conditions and standard deviations in mesocosm chambers determined from discrete seawater samples collected across six sampling time points during the challenge experiment. Each treatment was carried out in five replicate tanks, and observations were taken across six time points for each variable, resulting in a total of 30 measurements per variable (n = 30). TCO_2_ stands for the concentration of total carbon dioxide. The variables Ω_Ar_ and Ω_Ca_ indicate the saturation states of Aragonite and Calcite, respectively. The term pH_T_ represents the total scale pH, which accounts for all acid species within the sampled seawater.

| Treatment | Temperature (°C) | Salinity (PSU) | TCO<sub>2</sub> (μmol kg<sup>-1</sup>) | Total alkalinity (μmol kg<sup>-1</sup>) | pCO<sub>2</sub> (μatm) | pH<sub>T</sub> | $\Omega_{Ar}$ | $\Omega_{Ca}$ |
| --- | --- | --- | --- | --- | --- | --- | --- | --- |
| 800μatm+18°C | 18.03 ± 0.31 | 28.95 ± 0.21 | 2024.45 ± 78.37 | 2121.01 ± 80.89 | 780.67 ± 192.80 | 7.79 ± 0.09 | 1.36 ± 0.24 | 2.13 ± 0.38 |
| 1600μatm+18°C | 18.14 ± 0.24 | 28.99 ± 0.13 | 2110.44 ± 73.09 | 2122.09 ± 75.72 | 1564.73 ± 150.33 | 7.50 ± 0.04 | 0.73 ± 0.08 | 1.14 ± 0.13 |
| 2800μatm+18°C | 18.17 ± 0.21 | 29.06 ± 0.13 | 2222.46 ± 88.09 | 2156.82 ± 90.09 | 2954.77 ± 268.08 | 7.24 ± 0.04 | 0.42 ± 0.05 | 0.66 ± 0.08 |
| 800μatm+24°C | 24.04 ± 0.49 | 29.1 ± 0.16 | 1941.62 ± 65.85 | 2056.89 ± 76.13 | 814.33 ± 158.92 | 7.76 ± 0.08 | 1.57 ± 0.27 | 2.42 ± 0.42 |
| 1600μatm+24°C | 24.11 ± 0.19 | 29.18 ± 0.14 | 2040.31 ± 93.40 | 2068.38 ± 98.75 | 1641.78 ± 175.29 | 7.48 ± 0.05 | 0.87 ± 0.11 | 1.34 ± 0.17 |
| 2800μatm+24°C | 23.94 ± 0.31 | 29.23 ± 0.14 | 2125.81 ± 58.66 | 2095.03 ± 55.02 | 2690.24 ± 247.10 | 7.28 ± 0.04 | 0.56 ± 0.05 | 0.87 ± 0.07 |

**Table S2.**
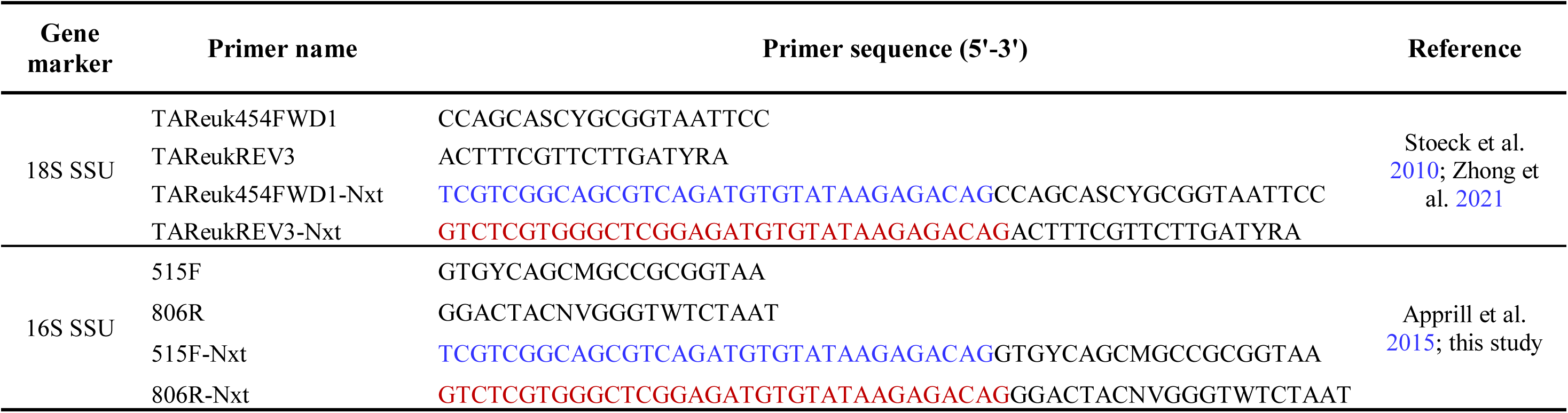
List of primers used in this study. Blue and red represent the overhanging adapter sequences for the forward and reverse primers, respectively.

**Table S3.** Dissimilarity of Pacific oyster spat associated prokaryotic and microeukaryotic community composition between two temperatures (18℃, 24℃) or among three *p*CO_2_ conditions (800, 1600, and 2800 µatm) across time revealed using permutational multivariate ANOVA (PERMANOVA) in the challenge experiment.

| Target | Community compositional dissimilarity | Time points | PERMANOVA r square and p value | Significance ($p \leq 0.05$: *, $p \leq 0.01$: **, $p \leq 0.001$: ***) |
| --- | --- | --- | --- | --- |
| Prokaryotic 16S rRNA gene | Between temperature | T1 to T6 together | $r^2 = 0.022, p = 0.002$ | ** |
| | Between temperature | T1 | $r^2 = 0.065, p = 0.264$ | |
| | Between temperature | T2 | $r^2 = 0.108, p = 0.001$ | *** |
| | Between temperature | T3 | $r^2 = 0.111, p = 0.001$ | *** |
| | Between temperature | T4 | $r^2 = 0.092, p = 0.02$ | * |
| | Between temperature | T5 | $r^2 = 0.127, p = 0.009$ | ** |
| | Between temperature | T6 | $r^2 = 0.123, p = 0.001$ | *** |
| | Among pCO<sub>2</sub> conditions | T1 to T6 together | $r^2 = 0.019, p = 0.386$ | |
| | Among pCO<sub>2</sub> conditions | T1 | $r^2 = 0.117, p = 0.492$ | |
| | Among pCO<sub>2</sub> conditions | T2 | $r^2 = 0.099, p = 0.905$ | |
| | Among pCO<sub>2</sub> conditions | T3 | $r^2 = 0.094, p = 0.938$ | |
| | Among pCO<sub>2</sub> conditions | T4 | $r^2 = 0.118, p = 0.445$ | |
| | Among pCO<sub>2</sub> conditions | T5 | $r^2 = 0.113, p = 0.496$ | |
| | Among pCO<sub>2</sub> conditions | T6 | $r^2 = 0.151, p = 0.124$ | |
| Microeukaryotic 18S rRNA gene | Between temperature | T1 to T6 together | $r^2 = 0.023, p = 0.002$ | ** |
| | Between temperature | T1 | $r^2 = 0.086, p = 0.055$ | |
| | Between temperature | T2 | $r^2 = 0.137, p = 0.001$ | *** |
| | Between temperature | T3 | $r^2 = 0.169, p = 0.001$ | *** |
| | Between temperature | T4 | $r^2 = 0.125, p = 0.002$ | ** |
| | Between temperature | T5 | $r^2 = 0.252, p = 0.001$ | *** |
| | Between temperature | T6 | $r^2 = 0.249, p = 0.001$ | *** |
| | Among pCO<sub>2</sub> conditions | T1 to T6 together | $r^2 = 0.018, p = 0.523$ | |
| | Among pCO<sub>2</sub> conditions | T1 | $r^2 = 0.116, p = 0.479$ | |
| | Among pCO<sub>2</sub> conditions | T2 | $r^2 = 0.106, p = 0.789$ | |
| | Among pCO<sub>2</sub> conditions | T3 | $r^2 = 0.101, p = 0.847$ | |
| | Among pCO<sub>2</sub> conditions | T4 | $r^2 = 0.098, p = 0.884$ | |
| | Among pCO<sub>2</sub> conditions | T5 | $r^2 = 0.089, p = 0.901$ | |
| | Among pCO<sub>2</sub> conditions | T6 | $r^2 = 0.092, p = 0.876$ | |

**Fig. S1.**
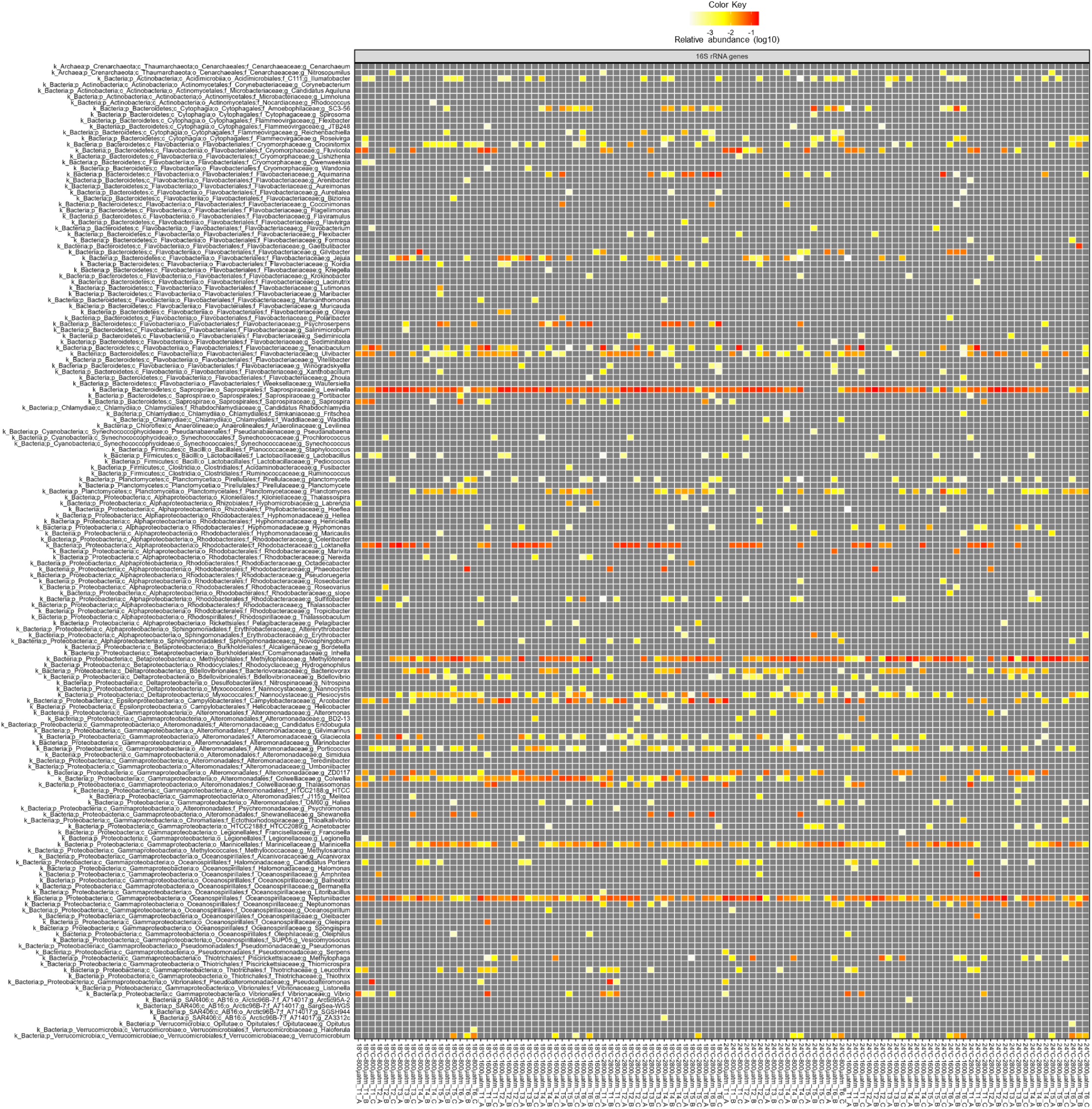
Genus-level relative abundance of prokaryotes in Pacific oyster spat samples across time and between different temperature and *p*CO_2_ conditions. Prokaryotes were inferred using 16S rRNA gene sequences of oyster samples. The color progression from grey to yellow to red represents the increasing relative abundance (log10) of 16S rRNA gene sequences.

**Fig. S2.**
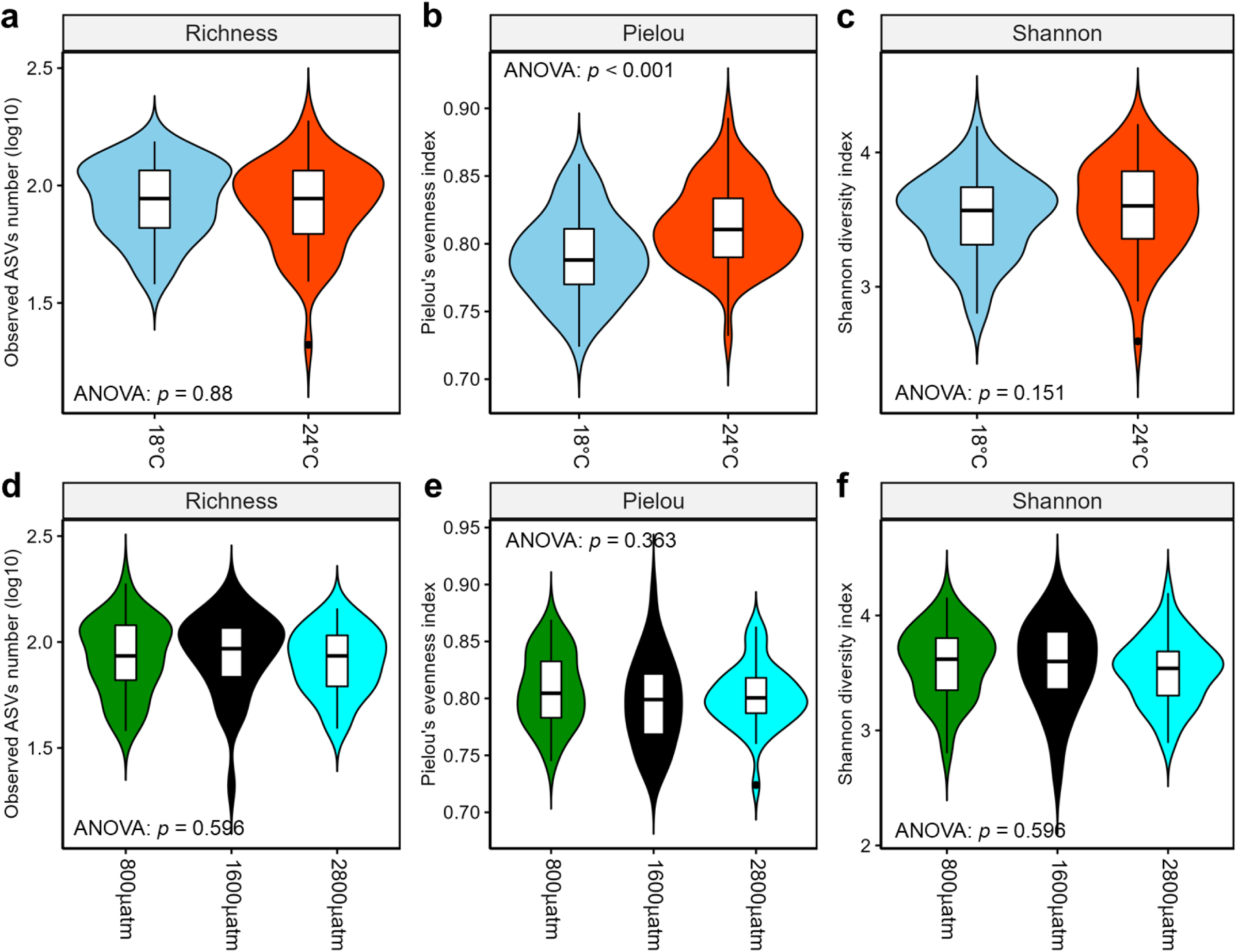
Alpha diversity of prokaryotic 16S rRNA genes across temperature and *p*CO_2_ treatments in Pacific oyster spat. The observed ASVs richness (**a, d**), Pielou’s *J* (**b, e**), and Shannon’s *H* (**c, f**) between two temperature (**a, b, c**) and among three *p*CO_2_ (**d, e, f**) treatments. The color sky-blue and red indicate temperatures of 18℃ and 24℃, respectively. Green, black, and cyan colors represent *p*CO_2_ levels at 800, 1600, and 2800 *µ*atm, respectively. The violin plot is a smoothed density plot of the data, with the violin’s width representing the frequency of observations in that range of values. The line in the middle of the violin plot represents the data’s median, and the square indicates the data’s mean.

**Fig. S3.**
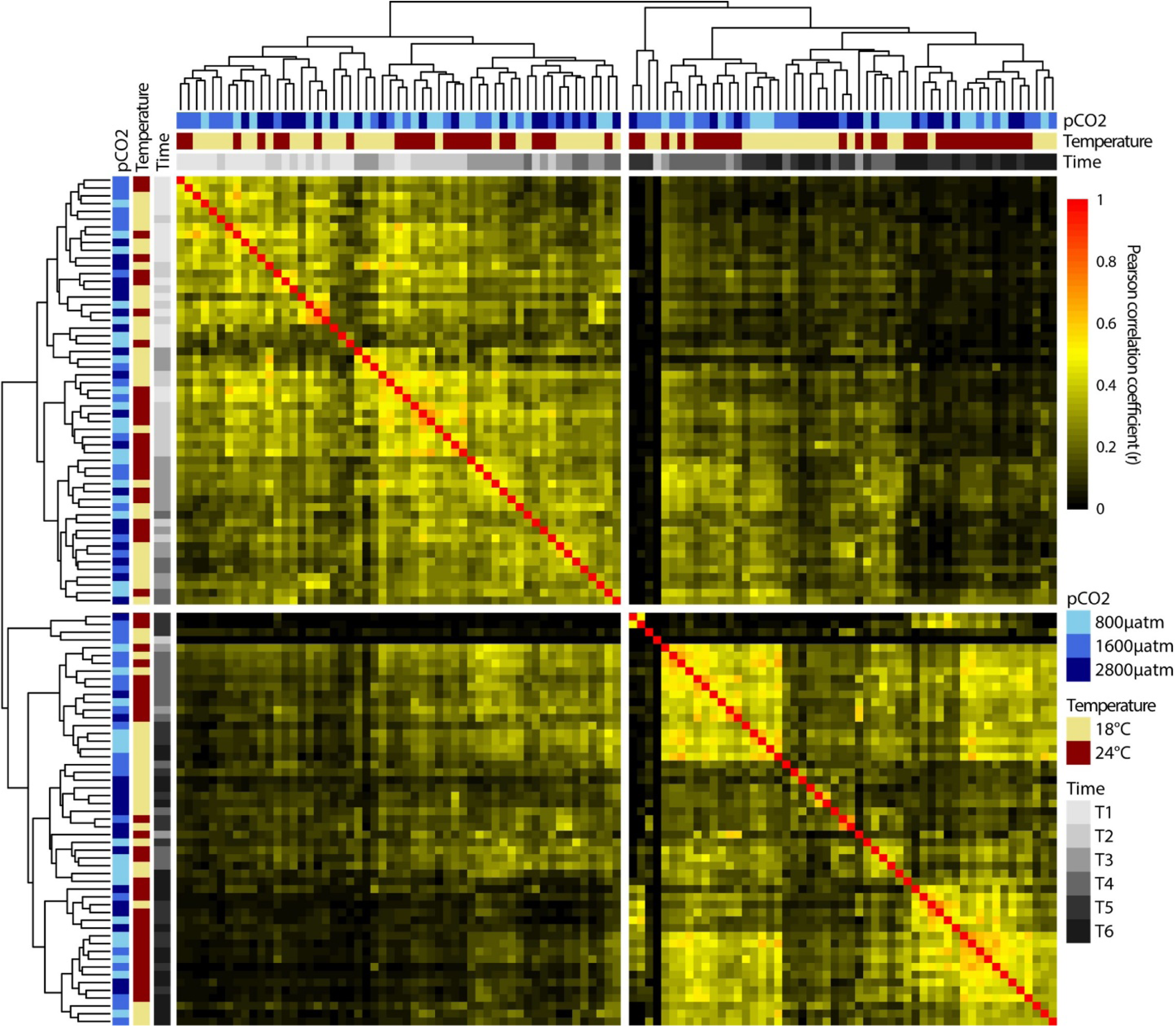
Pearson correlation coefficients between prokaryotic communities of Pacific oyster spat across time and treatments. Dendrogram to the left and top of the heatmap is clustered based on Bray-Curtis dissimilarity in the ordination of the Pearson correlation coefficients.

**Fig. S4.**
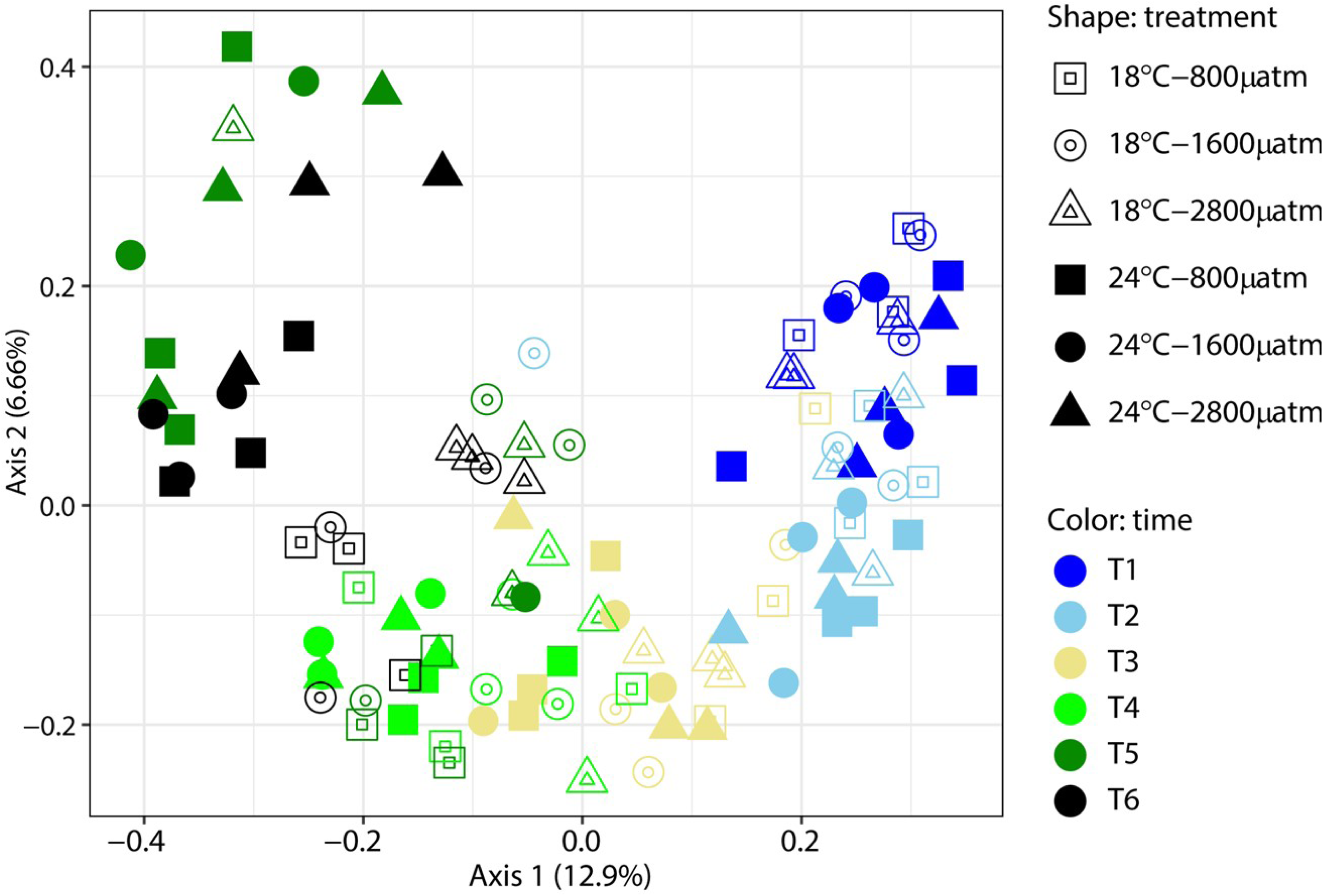
Principal Coordinate Analyses (PCoA) of the community structure of the prokaryotic 16S rRNA gene sequences in oyster spat samples subjected to different temperature and *p*CO2 treatments across time. The PCoA ordinates the weighted Unifrac distance metrics that are based on the presence/absence and the relative abundance of prokaryotic 16S rRNA gene ASVs. Symbol colours represent the time points of incubation, and the shapes indicate the treatment conditions.

**Fig. S5.**
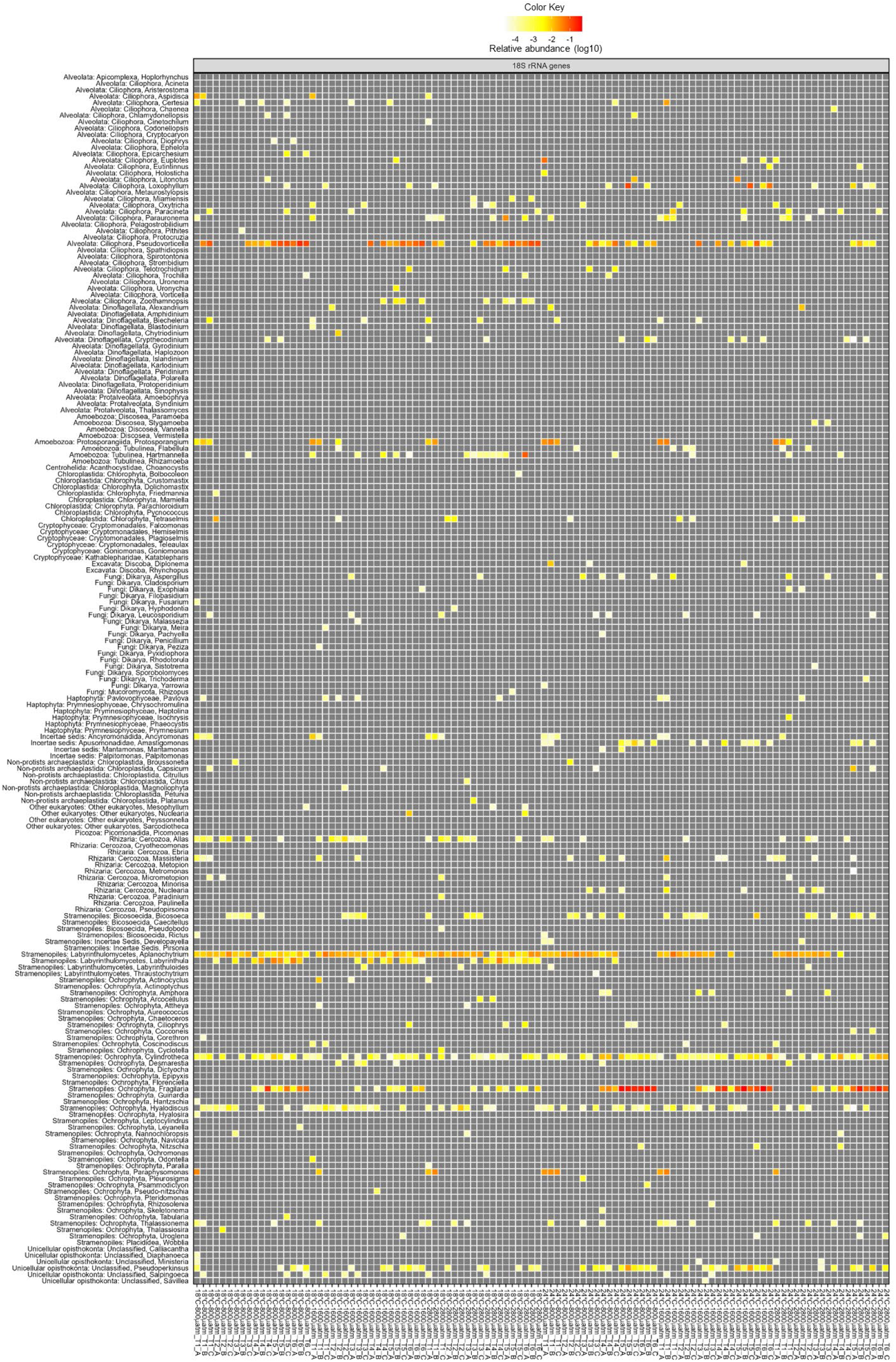
Genus-level relative abundance of microeukaryotes in Pacific oyster spat samples across time and between different temperature and *p*CO_2_ conditions. Microeukaryotes were inferred using 18S rRNA gene sequences of oyster samples. The colour progression from grey to yellow to red represents the increasing relative abundance (log10) of 18S rRNA gene sequences.

**Fig. S6.**
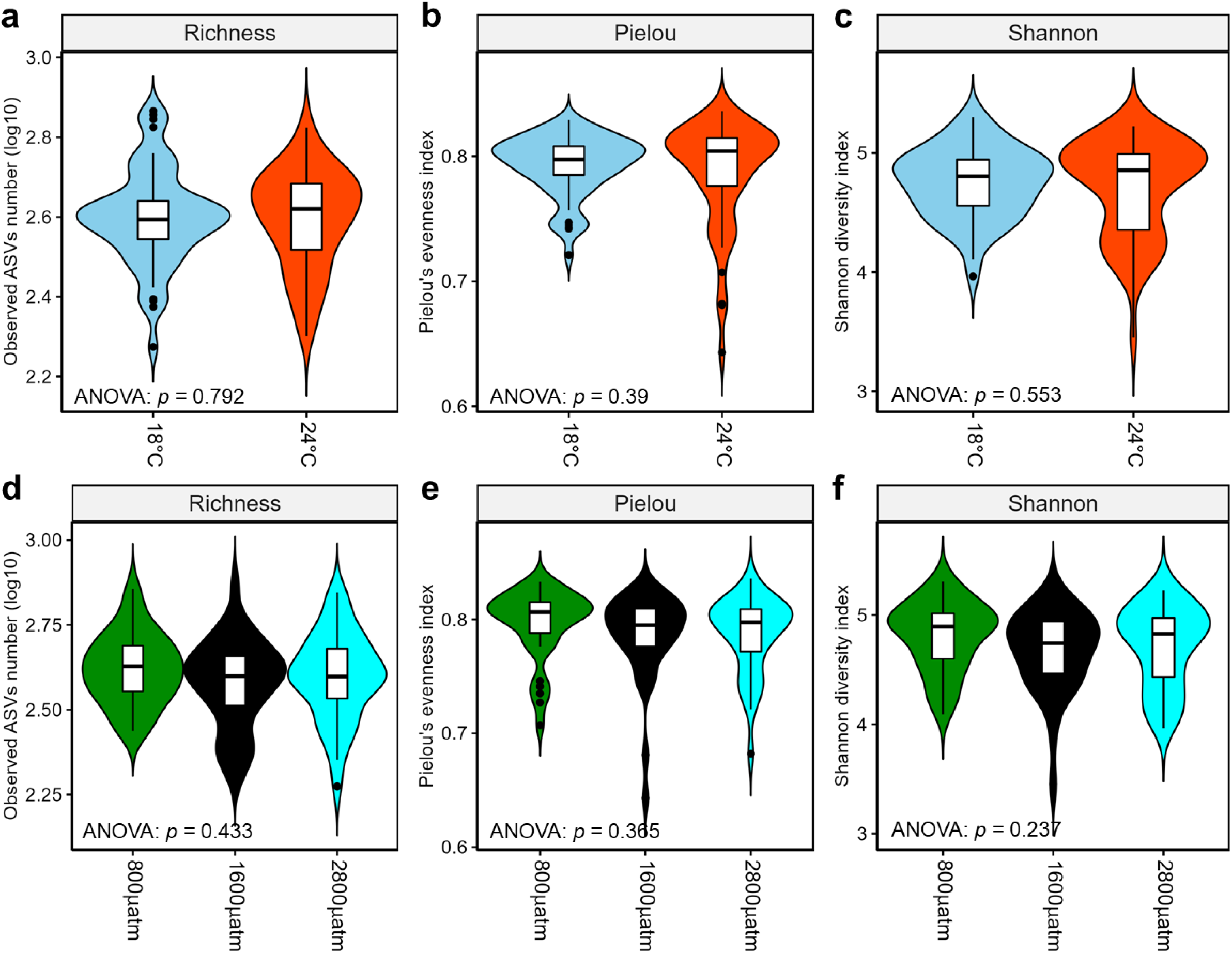
Alpha diversity of microeukaryotic 18S rRNA genes across temperature and *p*CO_2_ treatments in Pacific oyster spat. The observed ASVs richness (**a, d**), Pielou’s *J* (**b, e**), and Shannon’s *H* (**c, f**) between two temperature (**a, b, c**) and among three *p*CO_2_ (**d, e, f**) treatments. The color skyblue and red indicate temperatures of 18℃ and 24℃, respectively. Green, black, and cyan colors represent *p*CO_2_ levels at 800, 1600, and 2800 *µ*atm, respectively. The violin plot is a smoothed density plot of the data, with the violin’s width representing the frequency of observations in that range of values. The line in the middle of the violin plot represents the data’s median, and the square indicates the data’s mean.

**Fig. S7.**
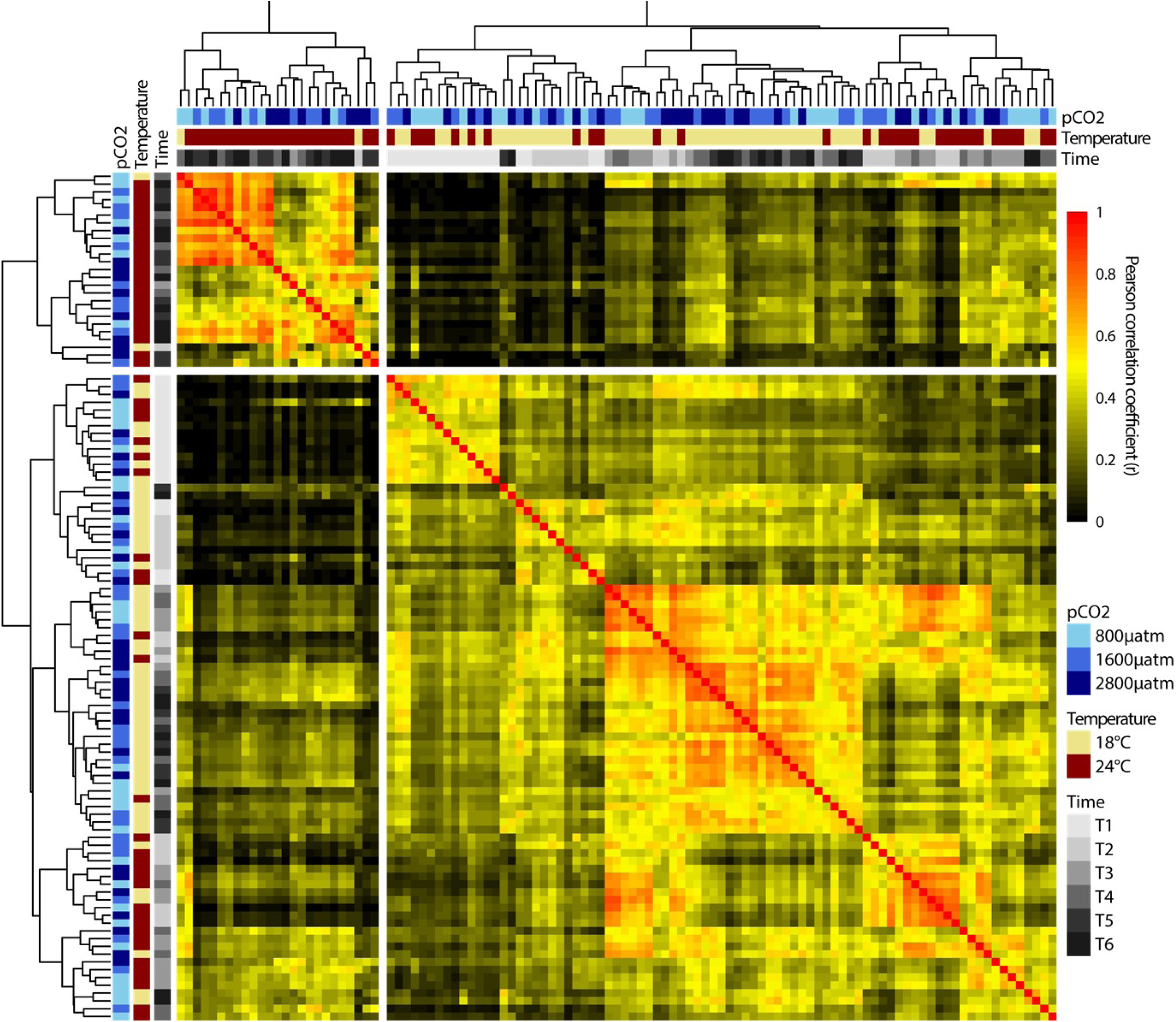
Pearson correlation coefficients between microeukaryotic communities of Pacific oyster spat across time and treatments. Dendrogram to the left and top of the heatmap is clustered based on Bray-Curtis dissimilarity in the ordination of the Pearson correlation coefficients.

**Fig. S8.**
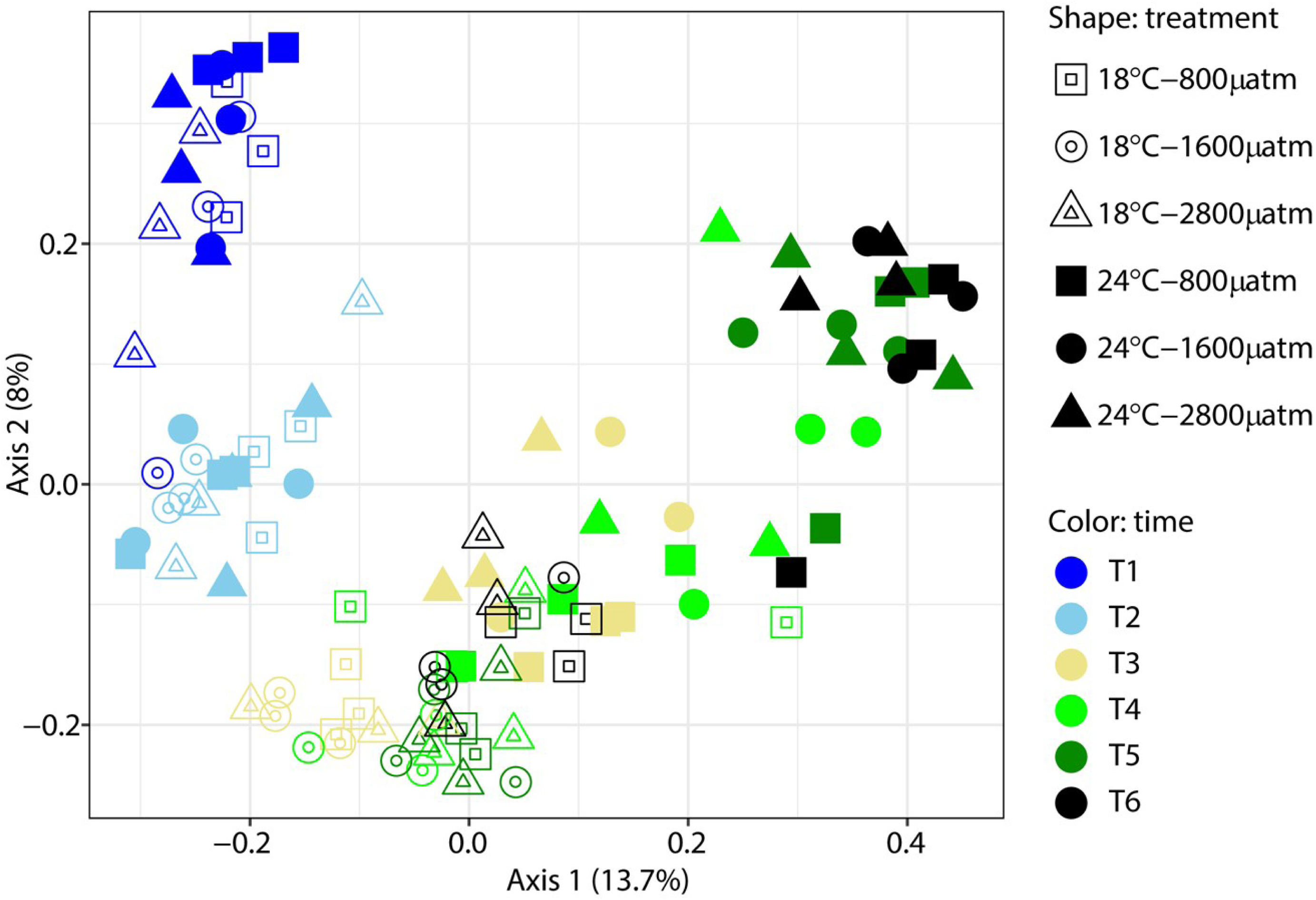
Principal Coordinate Analyses (PCoA) of the community structure of the microeukaryotic 18S rRNA gene sequences in oyster spat samples subjected to different temperature and *p*CO2 treatments across time. The PCoA ordinates the weighted Unifrac distance metrics that are based on the presence/absence and the relative abundance of microeukaryotic 18S rRNA gene ASVs. Symbol colours represent the time points of incubation, and the shapes indicate the treatment conditions.

**Fig. S9.**
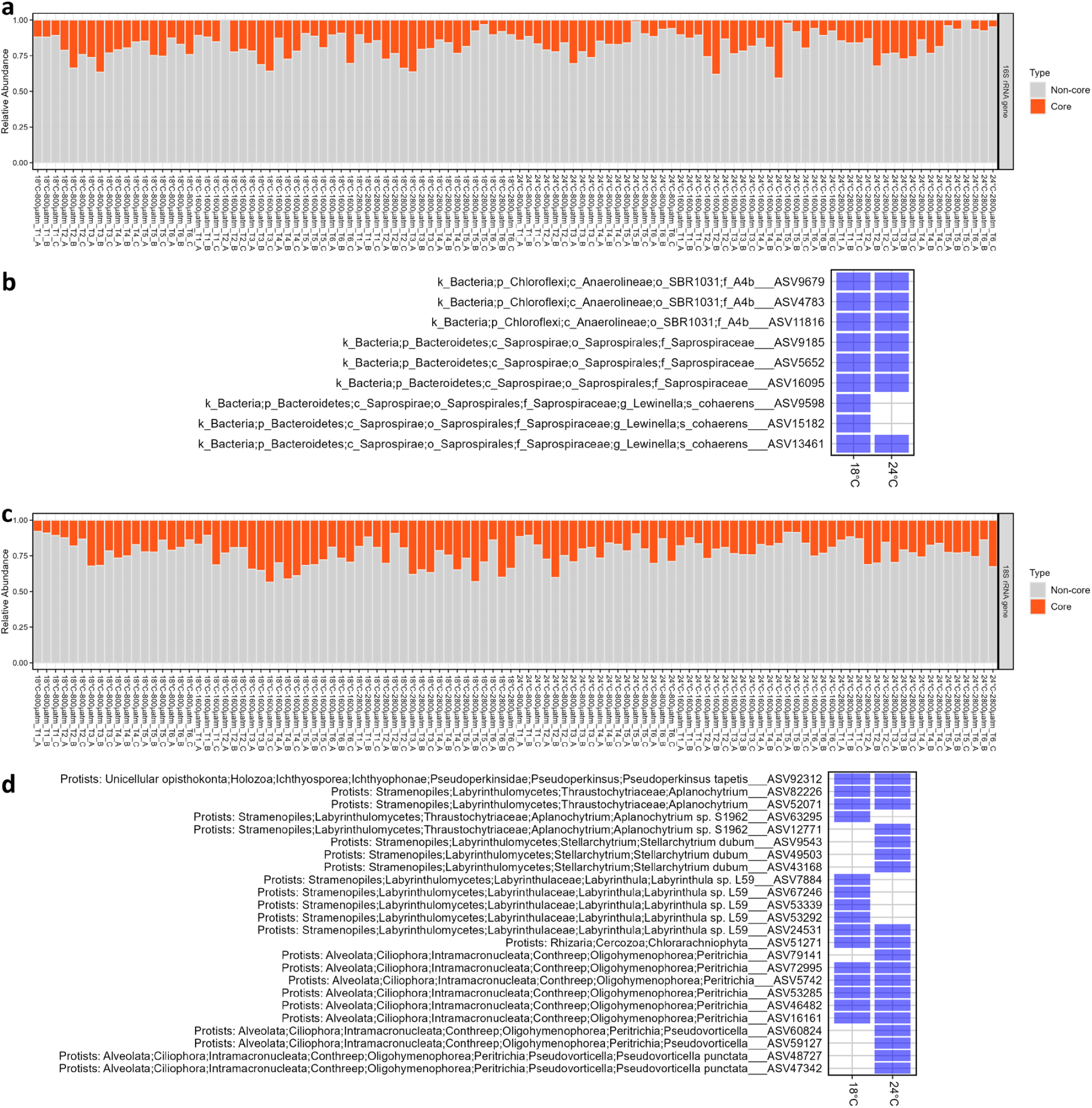
Core constituents of prokaryotic (a, b) and eukaryotic (c, d) microbiota in Pacific oyster spat samples across time in each temperature treatment. **a & c** Relative abundance of core and non-core constituents of the prokaryotic (**a**) and eukaryotic (**c**) microbiota in oyster spat samples of the challenge study. **b & d** Presence/absence of these core constituents of the prokaryotic (**b**) and eukaryotic (**d**) microbiota in oyster spat samples of each temperature treatment across time. Blue squares indicate the presence of the core taxa.

**Fig. S10.**
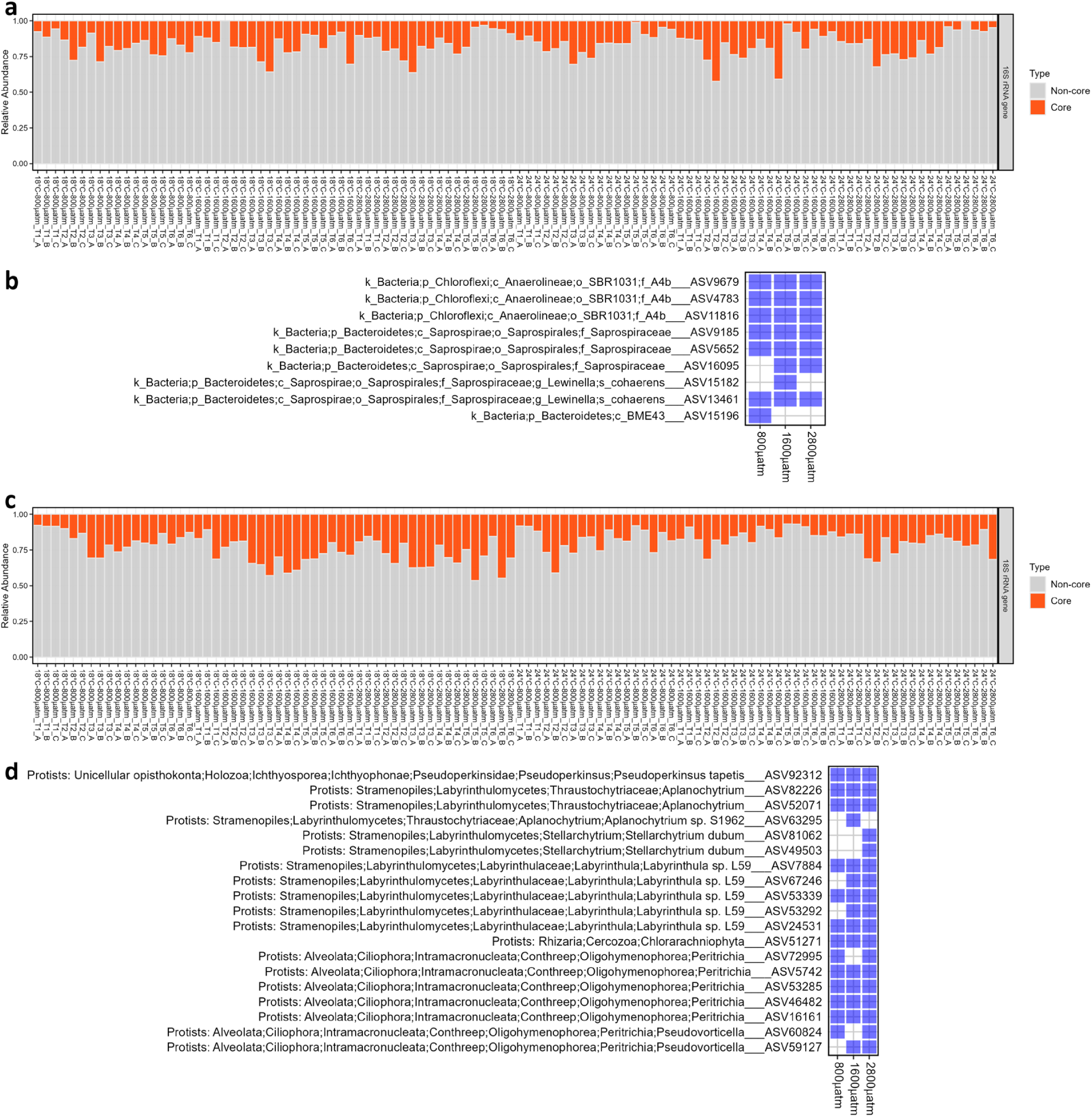
Core constituents of prokaryotic (a, b) and eukaryotic (c, d) microbiota in Pacific oyster spat samples across time in each *p*CO_2_ treatment. **a & c** Relative abundance of core and non-core constituents of the prokaryotic (**a**) and eukaryotic (**c**) microbiota in oyster spat samples of the challenge study. **b & d** Presence/absence of these core constituents of the prokaryotic (**b**) and eukaryotic (**d**) microbiota in oyster spat samples of each *p*CO_2_ treatment across time. Blue squares indicate the presence of the core taxa.

**Fig. S11.**
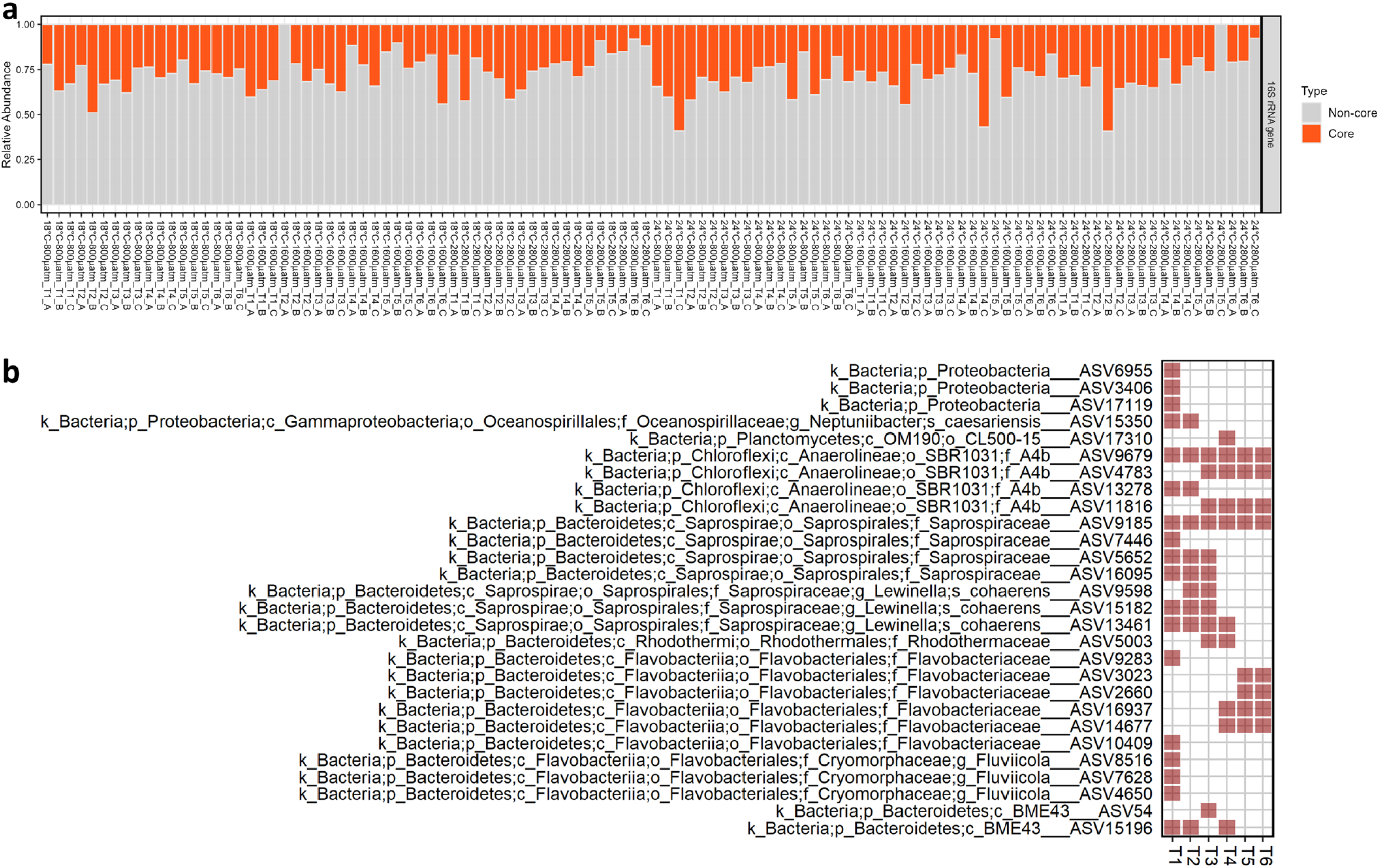
Core constituents of the Pacific oyster spat prokaryotic microbiota among different treatments at each time point. **a** Relative abundance of core and non-core constituents of the prokaryotic microbiota in oyster spat samples of the challenge study. **b** Presence/absence of these core constituents in oyster samples subjected to different treatments at each time point. The red squares indicate the presence of the taxa.

**Fig. S12.**
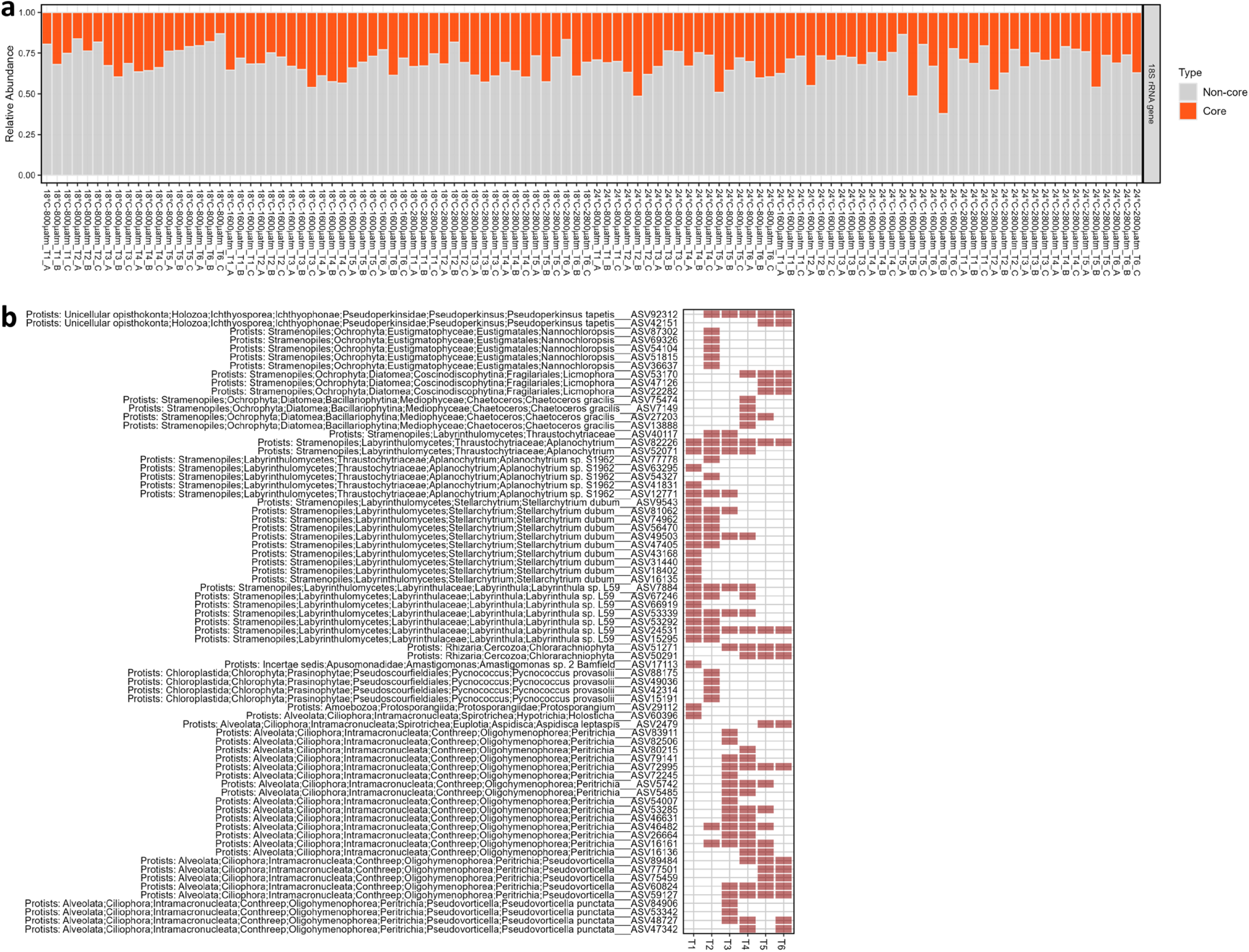
Core constituents of the Pacific oyster spat eukaryotic microbiota among different treatments at each time point. **a** Relative abundance of core and non-core constituents of the eukaryotic microbiota in oyster spat samples of the challenge study. **b** Presence/absence of these core constituents in oyster samples subjected to different treatments at each time point. The red squares indicate the presence of the taxa.

